# Exploring new dimensions of immune cell biology in *Anopheles gambiae* through genetic immunophenotyping

**DOI:** 10.1101/2024.10.22.619690

**Authors:** George-Rafael Samantsidis, Ryan C. Smith

## Abstract

Mosquito immune cells, or hemocytes, are integral components of the innate immune responses that define vector competence. However, the lack of genetic resources has limited their characterization and our understanding of their functional roles in immune signaling. To overcome these challenges, we engineered transgenic *Anopheles gambiae* that express fluorescent proteins under the control of candidate hemocyte promoters. Following the characterization of five transgenic constructs through gene expression and microscopy-based approaches, we examine mosquito immune cell populations by leveraging advanced spectral imaging flow cytometry. Our results comprehensively map the composition of mosquito hemocytes, classifying them into twelve distinct populations based on size, granularity, ploidy, phagocytic capacity, and the expression of PPO6, SPARC, and LRIM15 genetic markers. Together, our novel use of morphological properties and genetic markers provides increased resolution into our understanding of mosquito hemocytes, highlighting the complexity and plasticity of these immune cell populations, while providing the foundation for deeper investigations into their roles in immunity and pathogen transmission.

## Introduction

Immune cells are crucial components of the immune system in all Metazoa^1^, playing key roles in limiting infection, pathogen clearance, developmental regulation, and maintaining tissue homeostasis^2,3^. While vertebrate immune cells contribute to both innate and adaptive immune responses, invertebrates solely rely on innate immune mechanisms, where immune cells are essential to combat pathogen infections, maintain homeostasis, and ensure host survival^1,4^. Much of our understanding of insect cellular immunity has relied on studies in *Drosophila*^5,6^, where the genetically tractable system and extensive genetic resources have provided an important foundation for our understanding of cellular immune function and hematopoiesis in other insect systems. This includes mosquitoes, where comparable immune cell (hemocyte) subtypes have been described by morphological properties^7^.

This has led to the traditional classification of mosquito hemocytes into three major cell subtypes: granulocytes, oenocytoids, and prohemocytes^8,9^. Granulocytes, analogous to mammalian macrophages, are phagocytic cells with primary roles as immune sentinels to regulate immune homeostasis and pathogen elimination. Oenocytoids are most often implicated for their role in the production of prophenoloxidases (PPOs)^10^, while prohemocytes are presumed precursor cells thought to differentiate into the granulocyte and oenocytoid lineages^7,11^. However, our understanding of mosquito immune cell populations has primarily been limited to morphological observations which have led to discrepancies in cell classifications^12^ and numbers^8^. Previous studies using lectin-conjugated stains^7,13–15^ or lipophilic dyes^16–18^ have enhanced the visualization of mosquito immune cells, however evidence suggests that these methods provide general hemocyte staining and do not adequately resolve mosquito hemocyte subtypes^7,16^. Additionally, while the recent application of clodronate liposomes in mosquitoes has enabled techniques to examine the functional contributions of phagocytic granulocyte populations^11,19,20^, we still lack tools to evaluate the function of non-phagocytic cell types.

Immunophenotyping via flow cytometry has proven invaluable for studying the dynamics and heterogeneity of vertebrate immune cell populations^21,22^. In mosquitoes, flow cytometry has been used to identify phagocytic cell populations^23,24^ or in identifying changes in cell morphology and ploidy in response to blood-feeding^14,25^. However, further applications of flow cytometry in mosquitoes have been constrained by the lack of specific cell markers or antibodies needed to define mosquito immune cell subpopulations. Towards this goal, recent proteomic and single-cell studies have begun to address this limitation by identifying candidate hemocyte markers^11,26–28^, while expanding on the complexity of mosquito immune cell populations^11,28^.

The development of genetic resources in *Drosophila* has significantly advanced our understanding of insect immune cells and hematopoiesis, enabling precise labeling, manipulation of gene expression, and genetic ablation^29–33^. However, the development of similar tools in *Anopheles* has thus far been limited. Attempts to utilize the *Drosophila* hemocyte-specific hemolectin promoter in *Anopheles gambiae* have been met with mixed success, where a subset of hemocytes were fluorescently labeled only after blood-feeding^34^. An additional study using the *Anopheles* PPO6 promoter demonstrates the ability to label mosquito hemocyte populations^35^, which were later profiled in an initial scRNA study^36^, yet have not been thoroughly examined by their morphological characteristics to define their abundance and expression in hemocyte subtypes. Still, given the integral role of hemocytes in the immune responses and in modulating vector competence to virus^37,38^ and malaria parasite infection^13,15,19,39–41^, there remains a significant need to develop genetic tools to enhance our understanding of mosquito hemocyte function.

Here we examine an extended list of candidate hemocyte-specific promoters in *Anopheles gambiae*, identifying both pan-hemocyte and granulocyte-specific promoters that enhance our ability to visualize and characterize mosquito immune cells. Using these newly developed genetic resources, we performed imaging spectral flow cytometry to gain further insights into the mosquito hemocyte landscape. With these experiments enabling immune cell sorting based on physical and morphological properties, enhanced by the expression patterns of three genetic markers, we are able to classify mosquito immune cells with high-resolution. Additional experiments incorporating phagocytosis help to further resolve the phagocytic capacity of each cell population. In summary, our study provides a strong foundation for genetic immunophenotyping in mosquitoes, which offers significant potential to enhance our understanding of mosquito immune cell biology and mosquito-borne disease transmission.

## Results

### Establishment of pan-hemocyte and granulocyte-specific transgenic mosquitoes

With the aim of establishing transgenic *An. gambiae* that specifically express fluorescent markers in hemocytes, we used previously published single-cell^11^ and proteomic^26^ datasets for hemocytes to identify genes enriched across all hemocyte subtypes or specifically in granulocytes. As a result, we selected the promoter regions of NimB2 (AGAP029054), PPO6 (AGAP004977), and SPARC (AGAP000305) as putative panhemocyte promoters (**Fig. S1**) and the promoters of LRIM15 (AGAP007045) and SCRASP1 (AGAP005625) as putative granulocyte-specific promoters (**Fig. S1**). For each gene, genomic fragments including the 5’ UTR and ∼2000bp upstream of the putative transcription start site were used to incorporate the putative regulatory regions of each candidate gene promoter (**Additional File 1**). Five different piggyBac transposon constructs were generated containing the panhemocyte or granulocyte-specific promoters fused with CFP or GFP, respectively (**Fig. S1**). Each construct was successfully integrated into the *Anopheles* genome, as confirmed by splinkerette PCR, with at least two transgenic lines generated for each promoter construct (**Fig. S2**).

To determine potential position effects on promoter activity as a result from the random integration of piggyBac, we examined the expression levels of each gene marker across the different transgenic lines for each construct. We found significant differences in the three different lines generated with the PPO6-CFP construct, where the AP line displayed the highest expression. In contrast, the F1 line exhibited minimal CFP expression, and therefore, was not further evaluated (**Fig. S3**). While no significant differences were observed between lines for the SPARC-CFP, NimB2-CFP, or LRIM15-GFP constructs (**Fig. S3**), similar to PPO6-CFP, comparisons between the two SCRASP1-GFP lines displayed significantly higher transgene expression in the LGAP line relative to F2B line (**Fig. S3**). Of note, although *NimB2* was found to be significantly enriched in previous transcriptomic and proteomic datasets^11,26,42^, NimB2-CFP transgene expression was ∼10-30 times lower than that of the PPO6-CFP or SPARC-CFP constructs suggesting that the regulatory regions used for the construct may not be adequate to drive expression (**Fig. S3**).

### Molecular characterization of putative panhemocyte markers

Similar to previous studies with the PPO6 promoter^35^, we observed PPO6-driven CFP fluorescence in circulating hemocytes of whole mount mosquito larvae and pupae (**Fig. 1A)**. While we observe similar patterns of SPARC-driven CFP expression to that of PPO6 in larval and pupal hemocytes, the SPARC promoter also displayed visible activity in the fat body of both developmental stages (**Fig. 1B**). Additional qPCR analysis demonstrates that there are comparable levels of expression of the CFP marker in larvae and adults for both the PPO6 (**Fig. 1C**) and SPARC (**Fig. 1D**) promoters, with PPO6 promoter driving slightly higher levels of *CFP* expression in adults (**Fig. 1C**).

**Figure 1.**
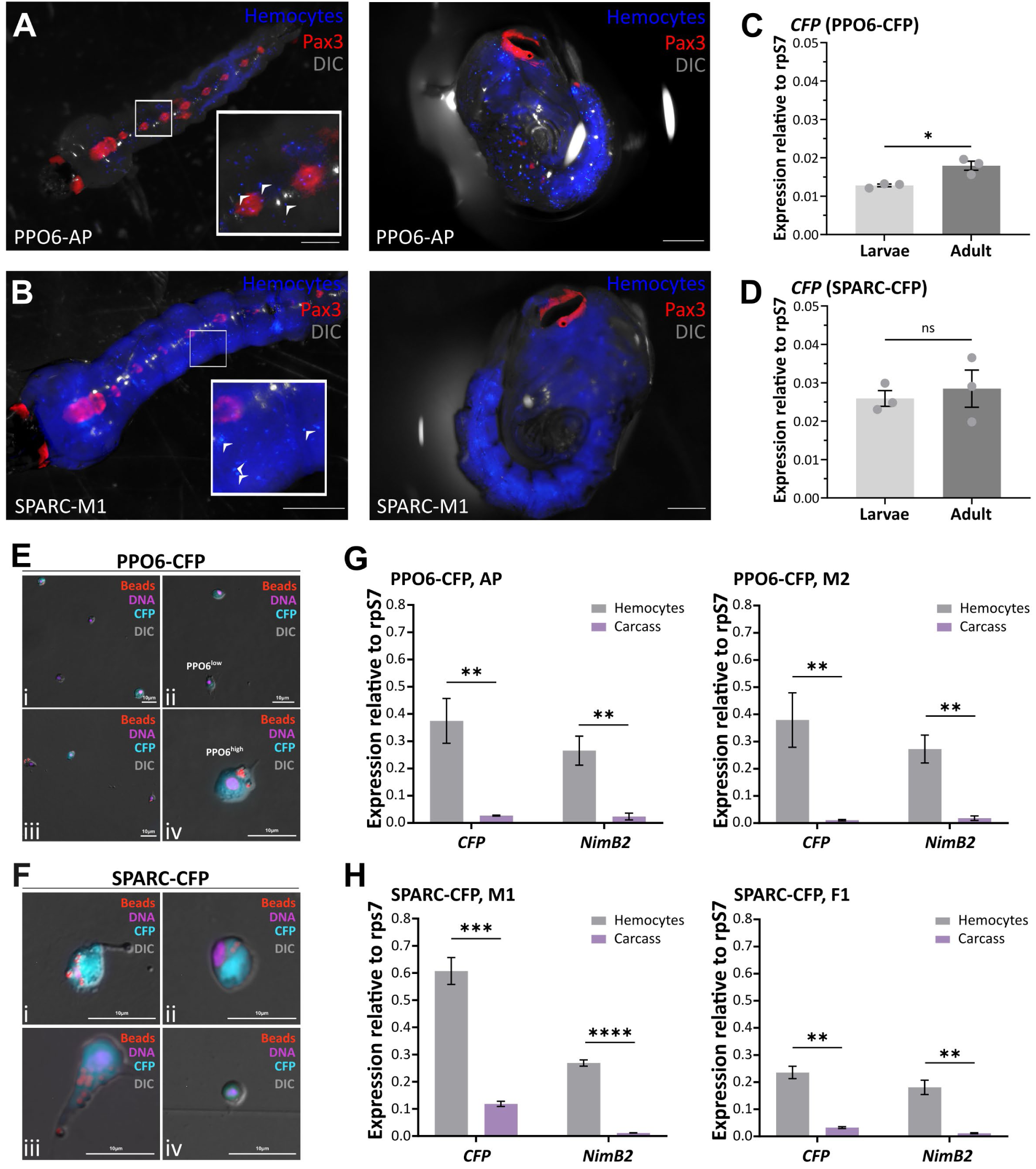
Molecular characterization of the putative panhemocyte markers PPO6 and SPARC. CFP fluorescence was examined in whole mount fourth-instar larvae and pupae from PPO6 (**A**) or SPARC (**B**) transgenic lines. Scale bars: 1mm. Potential differences in *CFP* expression between larvae and adult mosquitoes were examined by qPCR for both PPO6 (**C**) or SPARC (**D**) transgenic lines. *Ex vivo* analysis of mosquito hemocytes treated with beads indicates that PPO6^+^ (**E**) and SPARC^+^ (**F**) immune cell populations are comprised of phagocytic and non-phagocytic cells. Scale bars: 10μm. *CFP* expression is enriched in hemocytes compared to carcass as indicated by gene expression analysis in (**G**) PPO6 and (**H**) SPARC lines. The hemocyte-specific expression of *NimB2* was used as a positive control for gene expression analysis. Expression data are displayed relative to rpS7 expression, and bars represent the mean ± SE of three or four independent biological replicates. Significance was determined using multiple unpaired Student’s t-tests. Asterisks indicate significance (* *P* < 0.05, \*\**P* < 0.01, **** *P* < 0.0001). ns, not significant.

To further validate these findings, we perfused individual mosquitoes from each transgenic line to examine the fluorescence activity in adult hemocytes. Microscopic observations of hemocytes from PPO6-CFP adult transgenic mosquitoes revealed the existence of two distinct immune cell populations (**Fig. 1E**), referred to as PPO6^low^ and PPO6^high^, as previously described^19,35,36^. While most of the PPO6^+^ cells were classified as granulocytes, given their phagocytic capacity, none could be morphologically classified as other hemocyte subtypes (**Fig. 1E**). Similar observations were made for hemocytes perfused from SPARC-CFP mosquitoes, with varying CFP expression patterns detected among individual immune cells (**Fig. 1F**). Although the prevalence of SPARC^+^ cells (**Fig. 1F**) consisted primarily of phagocytic granulocytes with unique elongated projections extending outward from the cellular body, a limited number of cells displayed different morphological features, that were smaller in size and circular shaped with concentric nuclei potentially representative of oenocytoids or prohemocytes (**Fig. 1F**). Unfortunately, NimB2-CFP mosquitoes failed to display CFP fluorescence in perfused hemocytes (**Fig. S4**), consistent with the low expression levels of CFP in two different transgenic lines (**Fig. S3**). Additional experiments examining the potential that NimB2-CFP expression could be influenced by blood-feeding, as with previous hemocyte promoters in mosquitoes^34^, confirm the minimal levels of CFP expression under both naïve and blood-fed conditions (**Fig. S4**). Together, this confirms that the regulatory regions used for the NimB2 promoter are inadequate to drive heterologous expression with the promoter and was therefore not included in further analysis.

To examine the hemocyte-specificity of each promoter, we examined CFP expression in perfused hemolymph and carcass tissues for the PPO6-CFP and SPARC-CFP transgenic lines. Similar to the endogenous expression of the hemocyte-specific gene NimB2^11,19,36^, *CFP* expression was significantly enriched in hemocytes compared to carcass tissues for both the PPO6-CFP (**Fig. 1G**) and SPARC-CFP (**Fig. 1H**) transgenic constructs, with similar patterns of expression among different transgenic lines. However, the enrichment of *CFP* expression in hemocytes was less pronounced in SPARC-CFP mosquitoes when compared to PPO6-CFP transgenics (SPARC-CFP: ∼7X, PPO-CFP:25X, *p*= 0.0022), suggesting that CFP may be expressed in other mosquito tissues such as the fat body (**Fig. 1B**). Additional experiments using clodronate-liposomes to deplete phagocytic granulocyte populations^11,19,20^ further support the enrichment of PPO6-CFP, but not of SPARC-CFP, in granulocyte populations (**Fig. S5**). This lack of reduction in hemocyte-specific transgene expression could result from the activity of the promoter in non-phagocytic hemocytes that would be resistant to clodronate treatment or from leaky expression in other tissues beyond that of the immune cells, similar to what has been observed for the larval and pupal stages of SPARC-CFP mosquitoes (**Fig. 1B**).

### Molecular characterization of putative granulocyte markers

Granulocytes are central components of the mosquito innate immune responses that contribute to pathogen recognition and killing^19,40^. With previous studies featuring LRIM15 SCRASP1 prominently as granulocyte markers based on gene and protein expression analyses^11,26^, we opted to examine these gene promoters for their ability to drive granulocyte-specific expression. Unlike PPO6-CFP and SPARC-CFP transgenic lines, GFP fluorescence was not visibly detected in transgenic larvae of the LRIM15-GFP and SCRASP1-GFP lines (**Fig. 2A** and **2B**). Additional experiments using qPCR to compare the levels of *GFP* between larvae and adults for each transgenic line verified these observations and highlighted the specificity of these promoters to the adult stages (**Fig. 2C** and **2D**). Perfusion of LRIM15-GFP transgenic mosquitoes followed by immunostaining confirmed GFP expression in phagocytic granulocyte populations, consisting of cells with high and low patterns of GFP fluorescence that were observed in both native and fixed conditions (**Fig. 2E** and **Fig. S6**). In contrast with the LRIM15-GFP construct, SCRASP1-GFP hemocyte populations had much lower proportions of GFP-positive cells, with sizes ranging from approximately 3 to 10 microns (**Fig. 2F** and **Fig. S6**).

**Figure 2.**
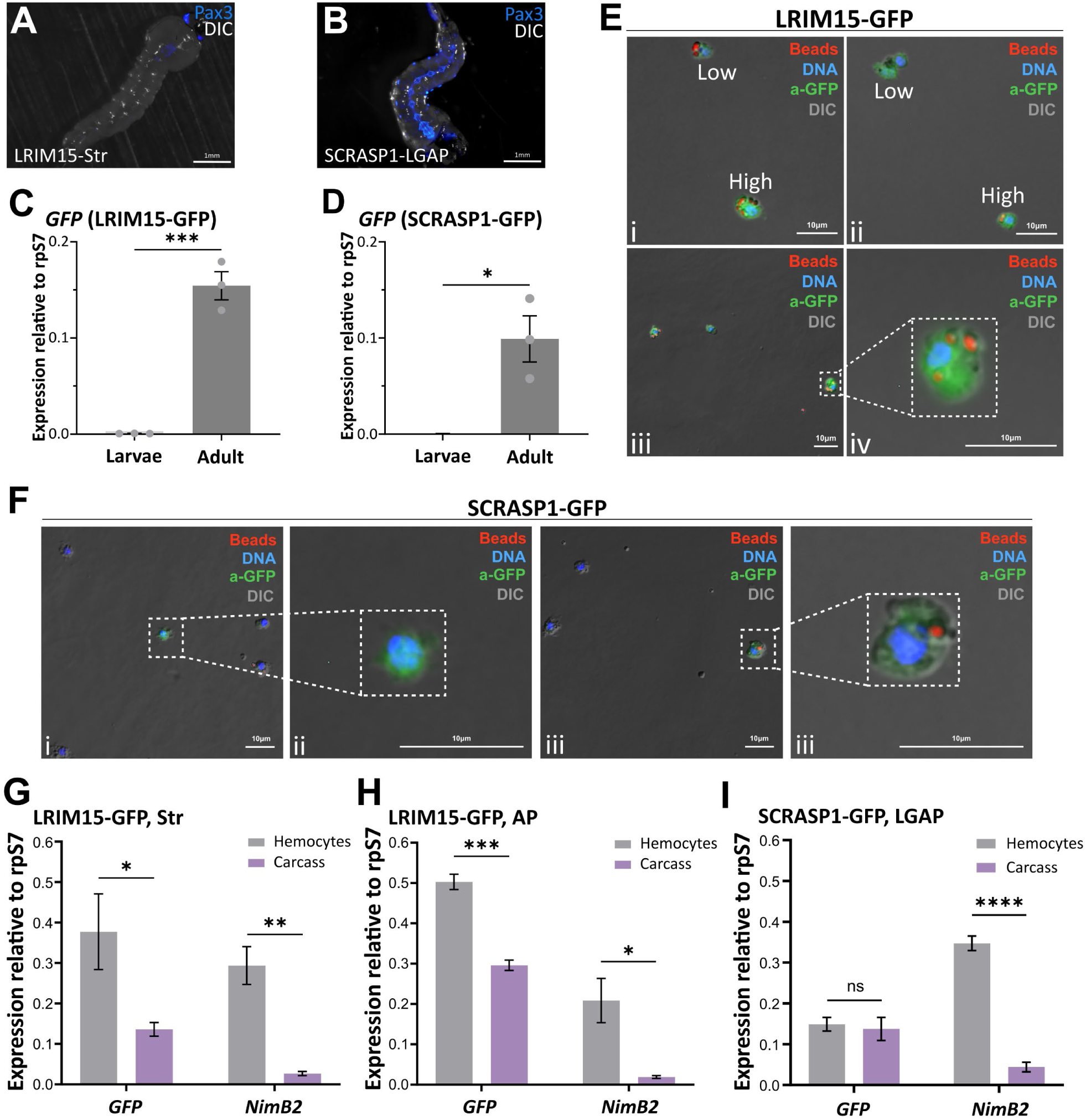
Molecular characterization of the putative granulocyte markers, LRIM15 and SCRASP1. GFP fluorescence was examined in whole mount fourth-instar larvae from LRIM15 (**A**) or SCRASP1 (**B**) transgenic lines. Scale bars: 1mm. Potential differences in *GFP* expression between larvae and adult mosquitoes were examined by qPCR for both LRIM15 (**C**) or SCRASP1 (**D**) transgenic lines. Immunostaining of adult hemocytes using an antibody specific to GFP (a-GFP) reveals various GFP^+^ populations with respect to fluorescence intensity and phagocytic capacity for (**E**) LRIM15-GFP and (**F**) SCRASP1-GFP transgenics. Hemocytes from the SCRASP1 line are comprised of cells with low GFP expression and size varying between 3-7μm (**F**). Scale bar: 10μm. *GFP* expression was enriched in hemocyte populations as compared to carcass tissue in both LRIM15 lines (**G** and **H**), although no difference was observed in the SCRASP1 mosquitoes (**I**). The hemocyte-specific expression of *NimB2* was used as a positive control for gene expression analysis. Expression data are displayed relative to rpS7 expression, and bars represent the mean ± SE of three to four independent biological replicates. Significance was determined using multiple unpaired Student’s t-tests. Asterisks indicate significance (* *P* < 0.05, \*\**P* < 0.01, **** *P* < 0.0001). ns, not significant.

The specificity of granulocyte-specific promoters was further evaluated by comparing *GFP* expression in perfused hemolymph with carcass tissues. While LRIM15 promoter activity was ∼3 times higher in hemocytes than in carcass tissues, there was no difference in GFP expression between tissues for the SCRASP1-GFP construct (**Fig. 2G-I**). To further confirm these observations, we again employed the use of clodronate liposomes to determine the effects of granulocyte depletion on *GFP* for each granulocyte promoter construct. While injections with clodronate-liposomes decreased *NimB2* expression by ∼60% in each strain, suggestive of phagocytic granulocyte depletion, clodronate treatment significantly reduced *GFP* expression in the LRIM15-GFP lines but not in the SCRASP1-GFP line (**Fig. S5**). We view this limited effect of clodronate treatment on *GFP* expression in the SCRASP1-GFP line as the result of low activity levels of the transgene, combined with the leaky expression in other tissues as indicated by qPCR (**Fig. 2I**). While we cannot exclude that there are low levels of expression in non-target tissues under the LRIM15 promoter, our results support that LRIM15 serves as a valuable granulocyte-specific marker.

### Hemocyte fluorescent markers reveal dynamic shifts in response to blood feeding

Previous studies have suggested that mosquito hemocytes are dynamic and undergo significant changes in response to blood-feeding^14,25,26,43^. For this reason, we examined the influence of blood-feeding on marker expression for each of our PPO6, SPARC, and LRIM15 hemocyte promoter constructs using microscopy and gene expression methods. Based on the low abundance of GFP^+^ cells and weak patterns of GFP expression (**Fig. 2** and **Fig. S5**), the SCRASP1 construct was not included in further analysis.

The abundance of PPO6-CFP^+^ hemocytes remained stable between sugar-fed and 24 hours post-blood feeding, representing ∼10% of total immune cells (**Fig. 3A**) and consistent with patterns of PPO6-driven *CFP* gene expression analysis (**Fig. S7**). However, at 48hrs post-feeding, PPO6^+^ cells displayed a small but significant increase in abundance (**Fig. 3A**), which can be attributed to an expansion in the proportions of PPO6^low^ populations (**Fig. S8**). In contrast, to the patterns observed in PPO6 immune cell populations, SPARC-CFP^+^ cells were more prevalent and displayed temporal oscillations in their abundance. We observed a significant increase in the proportions of CFP^+^ hemocytes from naive sugar-fed to 24hrs post-blood meal (∼45% to ∼63%), yet by 48hrs post-feeding, SPARC^+^ cell proportions significantly declined to ∼33% (**Fig. 3B**). Despite this variation in cell populations, no changes in SPARC-driven *CFP* expression were measured (**Fig. S7**). Of note, LRIM15-GFP^+^ cells displayed an inverse phenotype in response to blood-feeding, with a significant reduction of GFP^+^ cell proportions at 24hrs post-blood feeding, before reverting back to baseline levels (∼30% of cells) at 48hrs post-feeding (**Fig. 3C**). This is further supported by a corresponding decrease in GFP expression at 24hrs post-blood feeding (**Fig. S7**). Together, these data suggest that *An. gambiae* hemocyte populations are heterogeneous in nature and plasticity as they respond to physiological signals such as blood-feeding.

**Figure 3.**
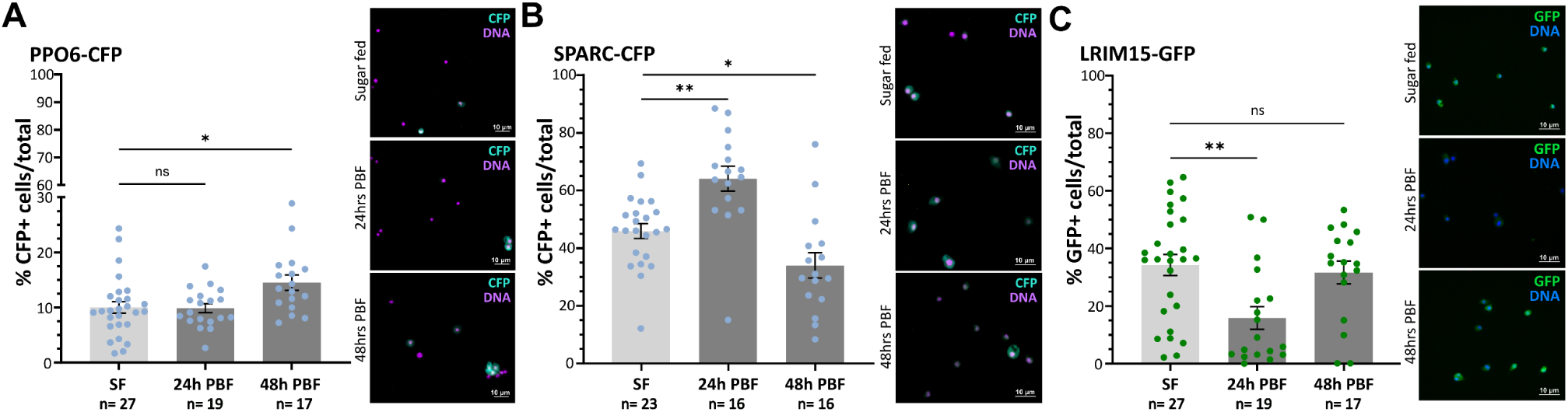
Blood-feeding influences PPO6^+^, LRIM15^+^, and SPARC^+^ immune cell populations. The percentage of PPO6^+^ (**A**), SPARC^+^ (**B**), and LRIM15^+^ **(C)** hemocytes were evaluated under sugar-fed (SF), at 24hrs post-feeding (24h PBF), and at 48hrs post-feeding. For each experimental condition, data from individual mosquitoes are displayed as dots, with no difference observed at 24hrs hemocyte populations increased at 24hrs post-blood meal and decreased at 48hrs. Conversely, hemocyte proportions displayed a significant decrease at 24hrs, which was recovered at 48hrs. For both A and B, the percentage of CFP+ or GFP+ cells of the total hemocytes are displayed as individuals (dots) and represented as the mean ± SE of three independent biological replicates. For each transgenic construct, representative images are displayed at right for each experimental condition. Statistical significance was determined by Mann-Whitney to compare the effects of blood-feeding at different time points. Asterisks indicate significance (\**P* < 0.05, \*\**P* < 0.01). ns, not significant; n numbers of individual mosquitoes examined. Scale bars represent 10μm.

### Mosquito immune cells comprise multiple subtypes based on ploidy and morphology

While conventional flow cytometry has been previously used to demonstrate differences in the DNA content of mosquito immune cells^11,14,25^, these studies have been limited by the lack of well-defined genetic markers and an inability to visualize cell heterogeneity in high resolution. With the advent of new technologies that combine spectral and imaging flow cytometry (IFC), and the development of the aforementioned genetic markers for PPO6^+^, SPARC^+^, and LRIM15^+^ immune cells, we now have the ability to examine mosquito immune cell populations at high resolution by combining analysis of cellular properties (DNA content, size, granularity) with morphological phenotypes (cell imaging).

To examine mosquito hemocyte populations by IFC, we first applied the use of this technology to characterize immune cells in wild-type *An. gambiae*. Using nuclear staining (DRAQ5) and real-time imaging, we gated *An. gambiae* hemocytes to select only for cells with clear morphology, whereas positive events for nuclear staining but without discernible cellular morphology, were gated out as debris (**Fig. S9**). Through this approach, mosquito hemocytes were most notably distinguished by DNA content or ploidy as previously^11,14,25^, resulting in the identification of five distinct subpopulations based on DNA content and designated as P1-5 (**Fig. 4A, Fig. S9**). Further examination of these P1-5 subpopulations using light loss (an indicator of cell size and granularity) enabled the separation of cells with similar ploidy into additional subgroups, ultimately resulting in the characterization of 12 immune cell subtypes displaying distinct cell properties of size and ploidy (**Fig. 4A, S10**). To better display the relationships between these immune cell subtypes, cells were visualized using UMAP and t-SNE (**Fig. 4B**). These results reveal clearly delineated cell clusters with minimal overlap, corroborating the efficiency of our gating strategy. Moreover, the clear separation of certain cell groups, including the P2.1, P4.1, P4.2, P5.1, and P5.2 groups (**Fig. 4B**), indicates the possibility of specialized functions for each of these immune cell subtypes. In contrast, the close spatial relationships observed for the remaining clusters potentially reflect their similar function and the possibility of phenotypic plasticity.

**Figure 4.**
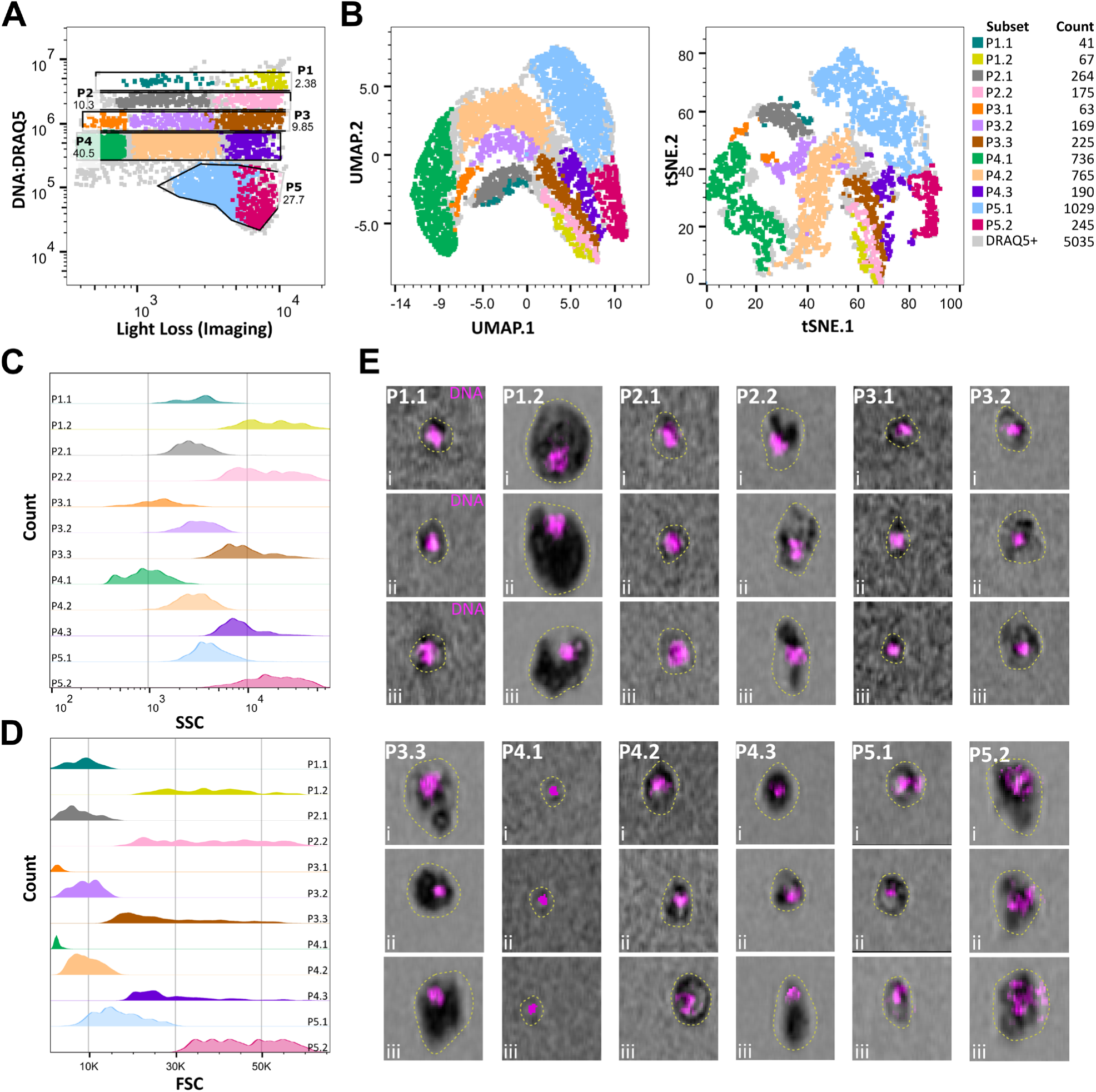
Imaging flow cytometry reveals multiple hemocyte subpopulations. The characterization of hemocytes from naïve, wild-type adult females enables the primary classification of hemocytes based on DRAQ5 signal (vertical y-axis) that revealed five cell clusters (**A**). When light loss measurements were accounted for (horizontal x-axis), additional subgroups were defined within each cell cluster (shaded by different colors). (**B**) Immune cell subtypes were analyzed by Uniform Manifold Approximation and Projection (UMAP) or t-Distributed Stochastic Neighbor Embedding (t-SNE) based on DRAQ5 signal, Maximum Intensity Forward Scatter (FSC), Maximum Intensity Side Scatter (SSC), and Maximum Intensity Light Loss characteristics, allowing for the clustering of hemocyte subpopulations according to relatedness. For each defined subtype, immune cell distributions are displayed for granularity (SSC; **C**) and size (FSC; **D**). **(E)** Representative images of each immune cell subcluster (P1.1-P5.2) are displayed, highlighting differences in size (outlined by dotted line), light loss, and DNA content (DRAQ5; magenta). UMAP analysis was performed using the Euclidean distance metric, and t-SNE was performed with opt-SNE learning configuration using FlowJo V10.10.0, including the following parameters: DRAQ5 signal, Maximum Intensity Forward Scatter (FSC), Maximum Intensity Side Scatter (SSC), and Maximum Intensity Light Loss. Each graph represents a single replicate of three independent biological experiments, with all data available in **Additional File 2**.

Granularity is one of the most prominent features of phagocytic immune cells across metazoa and has served as a defining feature of mosquito granulocytes. Side scatter analysis, a measurement of granularity, establish P3.1 and P4.1 clusters as the least granular cells, while the remaining clusters exhibited medium (clusters: P1.1, P2.1, P3.2, P4.2, and P5.1) or high (clusters: P1.2, P2.2, P3.3, P4.3, and P5.2) granularity (**Fig. 4C**). Forward scatter analysis, which allows for discrimination of cells by size, revealed a proportional relationship between granularity and cellular size, where those cells displaying the least granularity were the smallest in size, while clusters with medium or high granularity ranged from medium to large size (**Fig. 4D**). Among these, the P3.1 and P4.1 groups displayed the smallest size with minimal variation, while P1.2, P2.2, P3.3, P4.3 and P5.2 groups exhibited the largest variation in size (**Fig. 4D**). Real-time imaging corroborated these observations regarding the ploidy, size, and granularity of each cell cluster, providing additional insights into the morphological features and capturing the highly structured immune cell landscape of *An. gambiae* (**Fig. 4E**).

### Flow cytometry analysis of transgenic immune cell markers

Using our analysis of wild-type immune cell populations as a reference (**Fig. 4**), we next explored the properties of our SPARC^+^, PPO6^+^, and LRIM15^+^ immune cell populations to better understand the properties of the immune cell populations labeled by each genetic marker. Consistent with our microscopy analysis (**Fig. 3**), we see similar patterns of abundance for each transgenic construct, albeit at lower percentages of cells in our IFC analysis. Mosquitoes of the LRIM15-GFP line exhibited 15.2 ±2.5% of GFP^+^ cells (**Fig. 5A**), SPARC-CFP displayed the highest proportion with 27 ±1.02% of cells (**Fig. 5B**), and PPO6 labeled 8.7 ±1.02% of cells (**Fig. 5C**). Further analysis of fluorescent hemocytes in each transgenic line revealed their cellular composition (**Fig. 5D-I**). Hemocytes labeled by LRIM15-GFP were primarily composed of cells with medium to high granularity, with the P5.1 cluster accounting for 58.3% of the population (**Fig. 5D** and **5E**). SPARC-CFP hemocyte populations displayed strong similarity to LRIM15+ cells, with 61% of cells also belonging to the P5.1 cluster (**Fig. 5F** and **5G**). While PPO6-CFP fluorescent hemocytes were predominantly localized to cluster P5.1, they comprised a larger proportion of P2.2, P3.3, and P4.3 cells (**Fig. 5H** and **5I**). Together, these results indicate that while there are differences in the abundance of LRIM15^+^, SPARC^+^, and PPO6^+^ cells, their cellular properties (i.e. granularity) suggest that each of these cell subtypes are most likely granulocytes.

**Figure 5.**
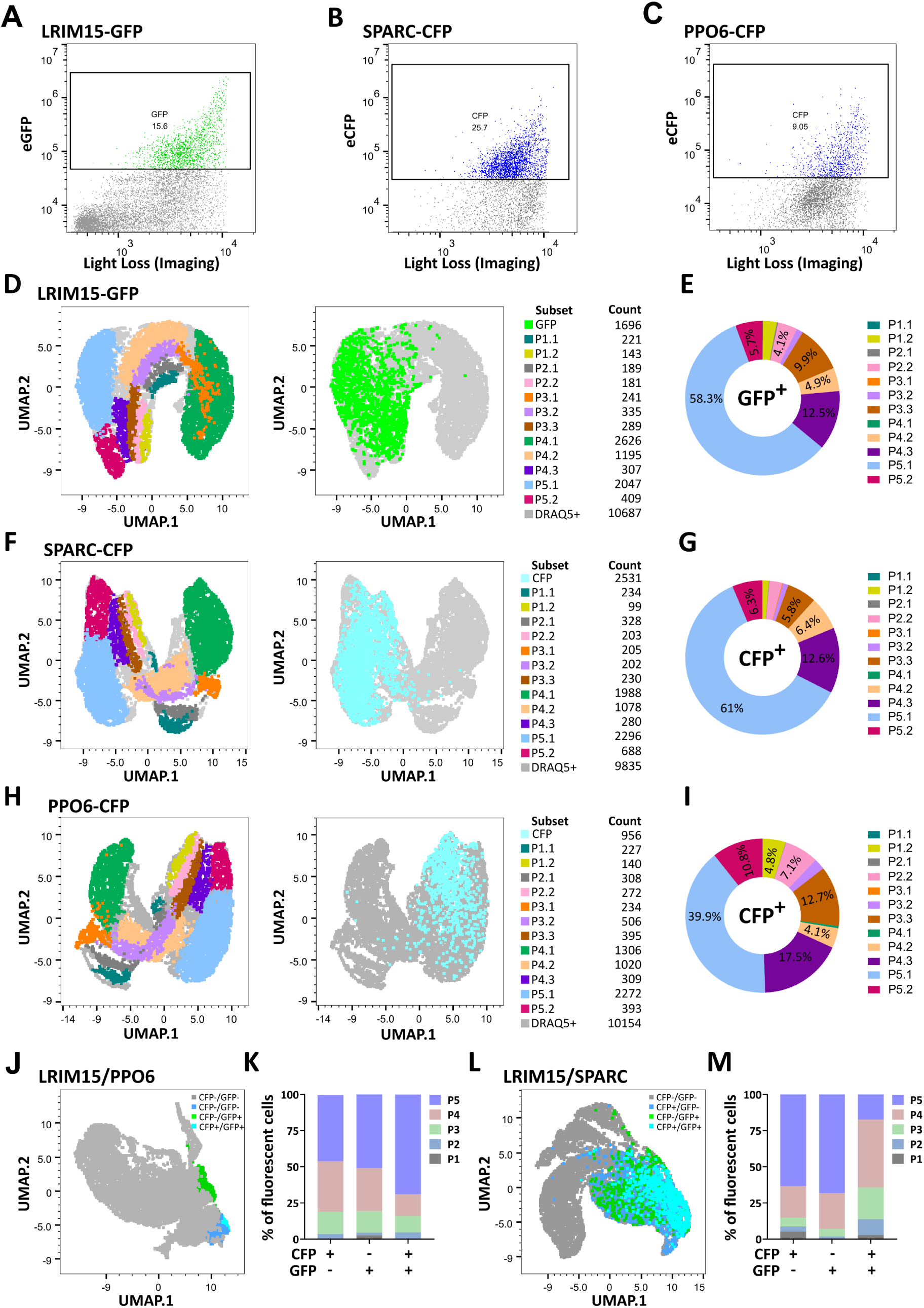
Flow cytometry analysis of LRIM15^+^, SPARC^+^, and PPO6^+^ hemocyte markers. Representative scatter plots of fluorescent hemocyte distribution in **(A)** LRIM15-GFP, **(B)** SPARC-CFP, and **(C)** PPO6-CFP mosquito lines. Cluster analysis using the UMAP dimensionality reduction technique was used to display fluorescent cell populations and to determine their composition using gating for each of the 12 hemocyte subpopulations identified in wild-type mosquitoes for LRIM15-GFP (**D, E**), SPARC-CFP (**F, G**), and PPO6-CFP constructs (**H, I**). To determine potential overlap between transgenic markers, crosses were performed to establish either LRIM15^+^/PPO6^+^ (**J**, **K**) or LRIM15^+^/SPARC^+^ (**L, M**) genetic backgrounds. For each background, the presence/absence of GFP^+^, CFP^+^, and GFP+/CFP+ cells were examined by overlaying fluorescent cell populations on the UMAP (**J**, **L**) or by hemocyte clusters (**K**, **M**). UMAP analysis was performed using the Euclidean distance metric using FlowJo V10.10.0, including the following parameters: DRAQ5 signal, Maximum Intensity Forward Scatter (FSC), Maximum Intensity Side Scatter (SSC), and Maximum Intensity Light Loss. Pie charts and graph bars were constructed using the average cell proportions of three biological replicates. Dot plots represent single replicates of three independent biological experiments, with all data available in **Additional File 2**.

Based on these similarities in the types of cells that are labeled with our respective LRIM15, SPARC, and PPO6 constructs, we wanted to examine the potential overlap between transgenic constructs. To address this, we outcrossed either SPARC-CFP or PPO6-CFP mosquitoes with LRIM15-GFP. Following the selection and establishment of mosquitoes with both fluorescent markers, we examined the presence/absence of CFP^+^ and GFP^+^ cells in these mixed genetic backgrounds. While there is some overlap between the PPO6 and LRIM15 markers (∼10% of total fluorescent cells), the two groups were clearly separated (**Fig. 5J** and **5K**), highlighting the distinct phenotypic properties and functions that distinguish between PPO6^+^ and LRIM15^+^ cells. In contrast, SPARC-CFP and LRIM15^+^ hemocytes displayed a higher degree of overlap with ∼30% of total fluorescent cells expressing both markers (**Fig. 5L**), with most CFP^+^/GFP^+^ cells belonging to the P4 cluster (**Fig. 5M**). In addition, while the P5 cluster individually represented >50% of the SPARC-CFP^+^ or LRIM15-GFP^+^ populations, it was underrepresented in cells co-expressing both genetic markers (**Fig. 5M**). Therefore, these data underscore the existence of distinct phenotypic features within morphologically similar immune cells and the further complexity of mosquito immune cell populations.

### Analysis of the phagocytic capacity of mosquito immune cell subtypes

In mosquitoes, granulocytes are the primary mediators of phagocytosis^44^, yet recent efforts have defined additional complexity in mosquito granulocyte populations^11,28^ and potential differences in the their phagocytic capacity^19^. To more closely examine mosquito immune cells involved in phagocytosis, we again employed IFC using fluorescent beads to examine the phagocytic capacity of cells.

Given the high fluorescence intensity of beads used in our experiments, we slightly adjusted our initial gating strategy to eliminate the potential spillover of red fluorescence into the DRAQ5^+^ channel (**Fig. S11**). The instrument’s high resolution, coupled with imaging, allowed us to exclude cell debris or bead singlets from the analysis to focus exclusively on phagocytic cells with stained nuclei (**Fig. S11**). When examined, phagocytic cells displayed high levels of light loss, suggesting that mosquito immune cells with greater density are more likely to engage in phagocytosis (**Fig. 6A**). Consistent with previous studies^19^, hemocytes displayed varying levels of phagocytic capacity indicative of the number of beads taken up by the cell, allowing us to classify them into three subpopulations: low, medium, and high (**Fig. 6B**). When our immune cell classifications are distinguished as either non-phagocytic or phagocytic cells, we see that some cell populations (P3.1 and P4.1) lack the ability to undergo phagocytosis, while others vary in their phagocytic capacity (**Fig. 6C** and **6D**). Those cells undergoing phagocytosis were primarily represented by P1.2, P2.2, P3.3, P4.3, P5.1, and P5.2 cell clusters (**Fig. 6C**), which exhibit larger size and greater light loss suggestive of granulocytes (**Fig. 6D**).

**Figure 6.**
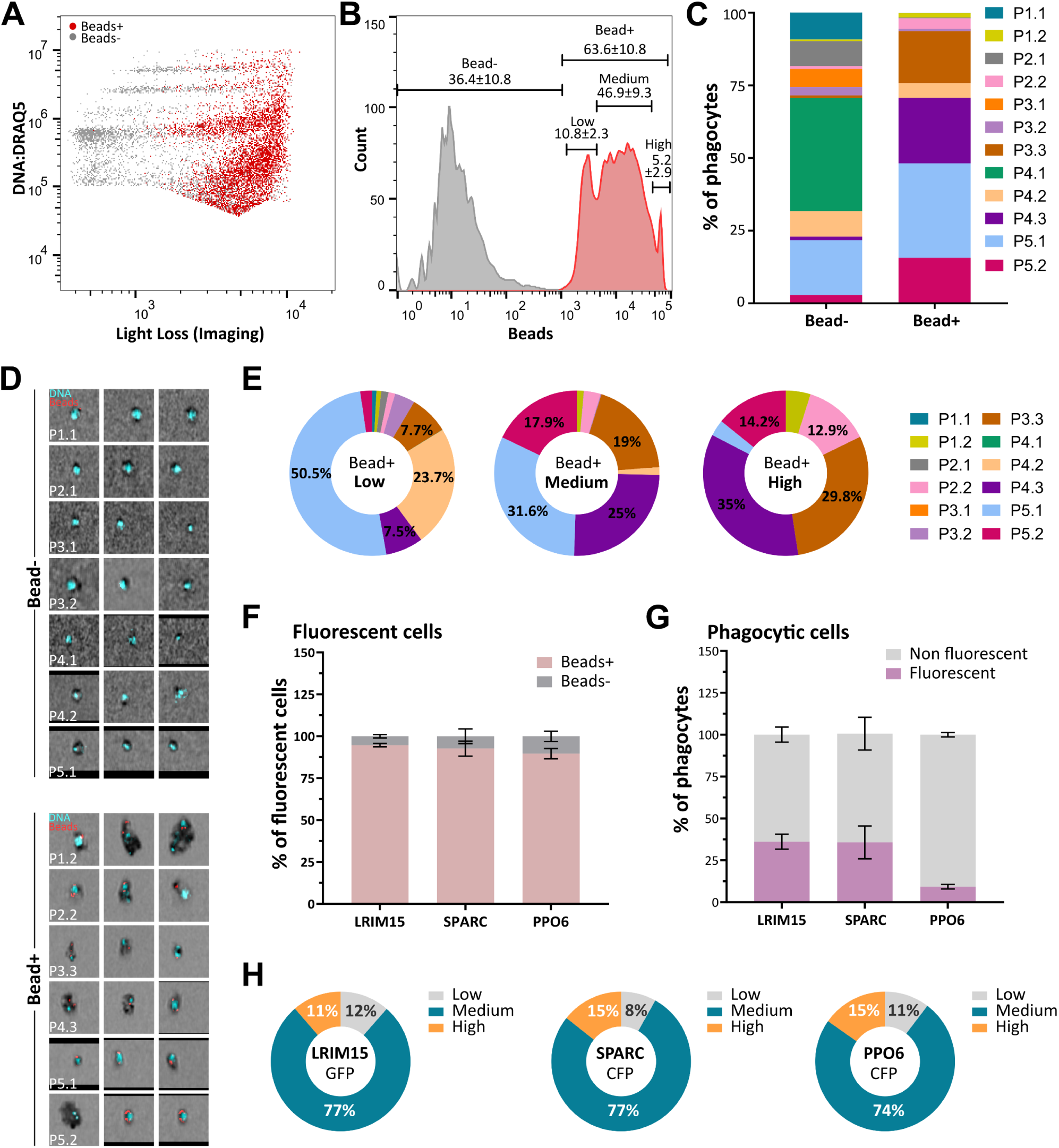
Mosquito immune cells vary in their phagocytic capacity. The injection of red fluorescent beads prior to perfusion enabled flow cytometry analysis of phagocytosis in mosquito immune cells (**A**) and the identification of multiple phagocytic immune cell phenotypes based on the intensity of bead signal (**B**). This resulted in the identification of non-phagocytic and phagocytic immune cell subtypes (**C**) that were confirmed by imaging (**D**). When phagocytic cells were further distinguished by bead signal intensity, we identified immune cell subtypes with low, medium, or high phagocytic capacity (**E**). Similar experiments with our LRIM15, SPARC, and PPO6 transgenic lines revealed the phagocytic ability of fluorescent immune cells (**F**) and the overall proportion of phagocytic immune cells (**G**) which are displayed as the mean ± SE of three biological replicates. (**H**) The phagocytic capacity of each transgenic line is visualized as the percentage of fluorescent cells displaying low, medium, or high bead-positive cells and displayed as the average of three independent biological replicates. Dot plots represent single replicates of three independent biological experiments, with all data available in **Additional File 3**.

To gain further insight into the observed differences in phagocytosis between immune cell clusters, we examined the composition of immune cells displaying low, medium, and high phagocytic capacity (**Fig. 6E**). Cells displaying “low” phagocytic capacity showed the highest representation of immune cell subtypes, while fewer immune cell clusters were represented in the “medium” and “high” phenotypes (**Fig. 6E**), suggesting that there is specialization of some immune cells to be more phagocytic. This is supported by the P5.1 cluster, which was enriched in cells with low and medium phagocytic capacity yet was underrepresented among cells displaying the highest bead uptake (**Fig. 6E**). Alternatively, these observations could also be partially justified by cell size, assuming a proportional relationship between cell size and phagocytic capacity. Consistent with this hypothesis, cells with the highest phagocytic capacity (P1.2, P2.2, P3.3, P4.3, and P5.2 clusters) (**Fig. 6E**) also were of the largest cell size and granularity among our immune cell subtypes (**Fig. 4C** and **4D**).

When we performed similar analysis with our LRIM15, SPARC, and PPO6 transgenic lines, >90% of fluorescent hemocytes displayed phagocytic capacity (**Fig. 6F**), providing further support for our observations that these transgenic constructs predominantly labeled populations of granulocytes (**Fig. 5**). However, only a subset of phagocytic cells was labeled in each of our transgenic lines, with LRIM15 and SPARC labeling up to 36% of phagocytes, while PPO6^+^ cells representing only ∼10% of phagocytic cells (**Fig. 6G**). When further examined for potential differences in phagocytosis, each of the transgenic markers labeled cell populations primarily comprised of cells with medium phagocytic capacity (**Fig. 6H**). This is consistent with the phagocytic abilities of P4.3 and P5.1 (**Fig. 6E**) which are predominantly labeled with each of the transgenic constructs (**Fig. 5**).

## Discussion

Hemocytes are integral components of mosquito innate immunity, with essential roles in defining vector competence and disease transmission. While mosquito hemocyte populations have traditionally been subdivided into three subtypes based on their morphological properties, recent studies have suggested a more complex and dynamic composition of immune cell subtypes^11,28^. However, the lack of genetic tools for mosquito hemocytes has been a significant hurdle for further studies to unravel their complexity, therefore causing a reliance on morphological properties that have only confounded their function^45^. Herein, we describe the development of multiple hemocyte markers and their utility to provide an unbiased classification of *A. gambiae* immune cells using genetic immunophenotyping.

With an initial goal to identify promoters that would comprehensively label all mosquito hemocyte populations or specifically target granulocytes, we identified three functional promoters (PPO6, SPARC, and LRIM15) able to successfully drive the robust expression of fluorescent markers in *A. gambiae* hemocytes. Additional promoter constructs using putative regulatory regions for SCRASP1 and NimB2 respectively displayed limited or no marker expression despite the prominence of both genes in previous studies of mosquito hemocytes^11,24,26,42^. This suggests that additional regulatory regions are likely required for both promoters to drive high levels of heterologous expression in mosquito immune cell populations. This may be addressed in the future through similar transposon-based experiments using extended regulatory regions or through a knock-in approach to drive expression using the endogenous gene^46^.

Based on previous single-cell studies^11,42^ and immunofluorescence experiments^24,25^, we had expected that the PPO6 and SPARC constructs could potentially be used to drive expression across hemocyte subtypes and serve as pan-hemocyte markers. This was supported by the previously established use of the PPO6 promoter to drive expression in *Anopheles* hemocytes^35,42^, although the abundance and cell distribution of PPO6^+^ cells has not previously been examined. Consistent with previous observations^35,42^, the PPO6 promoter drove hemocyte expression in both larval and adult stages at comparable levels. However, we found that the activity of the promoter was limited to a small subset of immune cells, accounting for only ∼10% hemocytes. This is in agreement with previous studies where the PPO6 promoter was active in only a subset of granulocytes^34^. Similar patterns of expression were observed for the SPARC-CFP construct in larvae and adults, although the proportions of SPARC^+^ hemocytes were significantly higher, reaching ∼50% in microscopy experiments and ∼25% of the total population via flow cytometry. However, these numbers fall short of achieving a “universal” promoter that would match the previous descriptions of PPO6 and SPARC expression. This may be attributed to increased mRNA stability of the transgene, which is negatively correlated with transcriptional rate^47,48^. Similar results have been observed in promoter characterization studies in lepidopteran species^49,50^, and may partially explain the limited activity of the promoters at levels below the limits of detection in non-labeled immune cell subtypes.

While SPARC expression is enriched in hemocytes, we also detected CFP expression in the fat body, which aligns with the tissue specificity of the *Drosophila* SPARC ortholog. Previous studies have shown that SPARC is localized in *Drosophila* hemocytes^51^ and fat body cells with a unique role in regulating the polymerization and deposition of collagen IV to sustain basal membrane integrity and fat body homeostasis^52–54^. Based on these similar expression patterns, this suggests that SPARC^+^ hemocytes may be involved in maintaining tissue homeostasis and production of the basal lamina, expanding the utility of the promoter beyond hemocyte function to other aspects of mosquito physiology.

With previous studies implicating the expression of LRIM15 with phagocytic immune cell populations^11,26^, the LRIM15 promoter construct performed as expected, driving strong fluorescent marker expression in mosquito granulocytes. However, we observed *GFP* expression and LRIM15^+^ cells only in adult mosquitoes, therefore suggesting that mosquito hemocytes undergo additional maturation or changes shortly after adult eclosion. Previous studies have highlighted differences between larval and adult immune responses, which includes an increase in phagocytic activity in adult mosquitoes^55^. While speculative, this suggests that the adult expression of LRIM15, and potentially other granulocyte-specific markers, may account for these immunological differences between mosquito life stages. Moreover, with only a limited understanding of larval hemocytes, these observations highlight the need for future studies to compare mosquito immune cell populations across development to better understand hematopoiesis and the stimuli that could influence immune maturation.

With evidence that mosquito hemocyte populations display plasticity and undergo significant alterations in response to physiological signals such as blood-feeding^14,25^, we provide initial proof-of-principle experiments that support that PPO6^+^, SPARC^+^, and LRIM15^+^ cells are dynamic in their abundance. For PPO6, we observe an increase in circulating PPO6^+^ cells at 48hrs post-blood feeding, with this change specifically attributed to the increased abundance of PPO6^low^ immune cell populations. We also observe an increase in SPARC^+^ cells at 24hrs post-blood meal, yet by 48hrs post-feeding there is a significant reduction in their abundance. In contrast, the abundance of LRIM15^+^ cells decreased at 24hrs post-blood meal and returned to normal levels by 48hrs post-feeding. While this is generally in agreement with previous observations that mosquito hemocytes undergo transient activation^25^, at present we still lack information as to how the physiological effects of blood-feeding or potentially other stimuli modify these mosquito immune cell populations. With each of the PPO6^+^, SPARC^+^, and LRIM15^+^ cells displaying properties of granulocytes, and only partial co-localization between these cell markers in our flow cytometry experiments, these data suggest that there is additional complexity in mosquito granulocyte populations that may reflect different levels of maturation, activation, or immune function.

While single-cell technologies have provided substantial resolution into the complexity of arthropod immune cells^11,28,51,56–58^, the further study of these immune cell populations in emerging model systems (such as mosquitoes and ticks) has been limited by the lack of cellular markers and genetic tools. Aided by the development of our PPO6^+^, SPARC^+^, and LRIM15^+^ transgenic lines and new advances in imaging flow cytometry, we used a multipronged approach to identify *An. gambiae* hemocyte populations and characterize their ploidy, size, granularity, morphology, and their phagocytic capacity. Consistent with previous studies^11,14,25^, our data demonstrate that mosquito hemocytes are readily distinguished by differences in DNA content or ploidy. While this may encompass some cells undergoing normal mitosis and cell division, the large proportion of immune cells displaying polyploidy is suggestive that endocycling (endomitosis or endoreplication) is an integral aspect of mosquito hemocyte biology. Cell polyploidy is common in insects and has been implicated in a variety of biological functions to increase transcriptional activity and protein secretion^59,60^. With previous studies in mosquito cell lines suggesting that endoreplication occurs in response to pathogen infection and is essential for immune priming^61,62^, polyploidy could represent a unique methodology used by mosquito immune cells for specialized immune functions or to enhance the response time to pathogen challenge.

When these aspects of ploidy are paired with traditional measurements of size and granularity which have been routinely used as a proxy of determining cell function^63,64^, we identify a total of twelve immune cell subpopulations in *An. gambiae*. Among these cell types, we see a clear delineation of non-phagocytic and phagocytic cells, which are readily distinguished by differences in size and granularity. With the advantage of our IFC methodology and the ability to visualize these cell types in addition to other physical measurements, we believe that the non-phagocytic cell types represent prohemocyte (clusters P3.1 and P4.1) and oenocytoid (P1.1 and P2.1) cell populations. In contrast, the phagocytic populations of immune cell are reminiscent of granulocytes (such as P3.3, P4.3, P5.1, and P5.2) and cells that likely correspond to the megacyte lineage (P1.2)^28,41^. Yet, given the observed differences in the phagocytic capacity of these cells, there appears to be significant complexity in these phagocytic cell populations. This is supported by the distinct patterns of PPO6^+^, SPARC^+^, and LRIM15^+^ cells within granulocyte populations that imply differences in immune maturation or specialized cell functions as previously suggested^11,28^. With the identification of additional immune cell markers and the expansion of our current genetic tools, we believe that future studies will be able to further delineate these mosquito immune cell subtypes and their contributions to mosquito innate immune function.

In summary, we believe that our study provides an essential foundation for future studies of mosquito immune cell biology where technical limitations have previously hindered progress. This includes the development of new genetic resources to enhance the visualization of hemocyte subtypes and the first demonstrated application of IFC technologies in an insect system that offer an increased resolution of mosquito immune cells. We believe that these important advancements now enable opportunities to address fundamental questions in mosquito hemocyte biology regarding hematopoiesis, cell differentiation, immune plasticity, and the cellular responses to a variety of physiological stimuli (blood-feeding, infection, etc.) using reproducible methodologies. Therefore, we believe these findings provide a critical resource for further investigations of mosquito hemocytes that will increase our knowledge and understanding of the integral roles of immune cell populations in mosquito vector competence.

## Materials and Methods

### Mosquito Rearing

Transgenic and wild-type *Anopheles gambiae* mosquitoes (Keele strain^66^) were reared at 27°C and 80% relative humidity, with a 14:10 hr light: dark photoperiod cycle. Larvae were fed on commercialized fish flakes (Tetramin, Tetra), while adults were maintained on a 10% sucrose solution and fed on commercial sheep blood (Hemostat) for egg production.

### Mosquito embryo transformation

Transgenic mosquitoes were generated using the piggyBac transposon system. *Anopheles gambiae* (Keele) preblastoderm embryos were injected by the Insect Transformation Facility at the University of Maryland Institute for Bioscience & Biotechnology Research. All injections were performed using an injection solution containing 150 ng/µL of piggyBac vector and 175 ng/µL of hyperactive piggyBac transposase mRNA^67–69^ under halocarbon oil as previously described^70^. After injections, the hatched insects that survived to adulthood were pooled based on sex and crossed with the wild-type strain *An. gambiae* (Keele). Progenies were screened for the expression of ECFP or DsRed integration markers at late larval stages. Individual transgenic lines were identified by distinct expression patterns of the ECFP or DsRed integration markers which were used to establish unique colonies.

### Hemocyte-specific transgenic mosquitoes

Hemocyte-specific *An. gambiae* reporter lines were generated by fusing the promoter sequences that drive universal hemocyte or granulocyte-specific gene expression to the fluorescent markers CFP and GFP, respectively. Using previous transcriptomic, proteomic, and functional data^11,71^, the genes NimB2 (AGAP029054), SPARC (AGAP000305) and PPO6 (AGAP004977) were selected as universal hemocyte markers, while LRIM15 (AGAP007045) and SCRASP1 (AGAP005625) were chosen as specific to granulocytes. Promoter sequences encompassing the 5’ untranslated regions and up to ∼2kb upstream of the Transcription Start Site (TSS) of each gene, were downloaded from Vectorbase and were either PCR amplified or underwent *de novo* synthesis (Integrated DNA Technologies, IDT) (**Additional File 1**). All primer pairs used for PCR amplification of the putative promoters are listed in **Table S1**. Following amplification, PCR products were initially subcloned in pJET1.2/blunt (Thermo Fisher) for sequence verification by Sanger sequencing (DNA Facility, Iowa State University) prior to cloning into the respective piggyBac constructs.

### Genomic DNA extraction

Genomic DNA was extracted from pools of ten adult mosquitoes as previously^72,73^ by homogenizing in Bender buffer (0.1M NaCl, 0.2M Sucrose, 0.1M Tris-HCl, 0.05M EDTA pH 9.1 and 0.5% SDS), followed by incubation at 65°C for 1 hr. After adding 15ul of 8M potassium acetate, samples were incubated for 45 min on ice and centrifuged for 10 min at maximum speed. Genomic DNA was ethanol-precipitated and resuspended in nuclease-free water.

### Plasmid construction

The open reading frame (ORF) of *DsRed* was excised from piggyBac-3xP3-DsRed^74^ with *NcoI-NotI* and replaced with either the *ECFP* ORF from piggyBac-ECFP-15xQUAS_TATA-mcd8-GFP-SV40^75^ (addgene: 104878) or *GFP* ORF amplified from an existing pJET1.2-T7-GFP plasmid^76^ with primers GFP-F-NcoI and GFP-R-NotI. Candidate hemocyte promoters were amplified with Phusion polymerase (ThermoFisher) using primers with *AscI* or *FseI-AsiSI* restriction sites respectively attached to the 5’-end of the forward or reverse primers **(Table S1)** and cloned into the *AscI* and *FseI* restriction sites of the desired piggyBac plasmid. The ORFs of CFP and GFP followed by SV40 termination sequence were inserted at the 3’ end of each candidate promoter using the restriction sites *AsiSI* and *FseI*. All plasmid sequences were confirmed by Sanger sequencing prior to microinjection, with sequences of each construct provided in **Additional File 1**.

### Mapping hemocyte-specific transgene insertion in *Anopheles* genome

To identify the integration sites of each hemocyte-specific transgene, we performed splinkerette PCR (spPCR) on the genomic DNA of each transgenic line as previously described^77,78^. Genomic DNA was extracted from pooled adult mosquitoes and digested with *BglII* or *MspI* for four hours. Splinkerette double-stranded oligos were synthesized to complement the sticky ends generated by *BglII* or *MspI*. Digested genomic DNA was ligated to the respective annealed splinkerette oligos with T4 DNA ligase (ThermoFisher) at 4°C overnight. PCR reactions were performed using Phusion polymerase (NEB) as previously described^78^. A list of all primers used for spPCR is summarized in **Table S2**. PCR fragments were gel purified using Gel DNA Recovery Kit (ZymoResearch) and cloned to pJET1.2/Blunt vector for Sanger sequencing. The recovered DNA sequences were mapped to *the An. gambiae PEST* reference genome using the blastn function in VectorBase.

### RNA extraction and gene expression analyses

Total RNA was extracted from whole mosquito samples using Trizol (Invitrogen, Carlsbad, CA). RNA samples prepared from perfused hemolymph samples were isolated using the Direct-Zol RNA miniprep kit (Zymo Research). Two micrograms of whole mosquito-derived or 200ng of hemolymph-derived total RNA were used for first-strand synthesis with the LunaScript RT SuperScript Kit (NEB). Gene expression analysis was performed with quantitative real-time PCR (qPCR) using PowerUp SYBRGreen Master Mix (Thermo Fisher Scientific). qPCR results were calculated using the 2^−ΔCt^ formula and normalized by subtracting the Ct values of the target genes from the Ct values of the internal reference, *rpS7*. All primers used for gene expression analyses are listed in **Table S3**.

### Transgene expression in response to blood-feeding

To determine the effects of blood-feeding on transgenic lines, adult transgenic mosquitoes (3-5 days old) were allowed to feed on defibrinated sheep blood for 5 min using an artificial membrane feeder. At 24 hrs post-blood feeding, engorged female mosquitoes were separated from unfed and used for RNA extraction and gene expression analysis. All blood-feeding experiments were repeated at least three times.

### Hemolymph perfusion

Mosquito adult hemolymph was collected by perfusion using an anticoagulant buffer of 60% v/v Schneider’s Insect medium, 10% v/v Fetal Bovine Serum, and 30% v/v citrate buffer (98 mM NaOH, 186 mM NaCl, 1.7 mM EDTA, and 41 mM citric acid; buffer pH 4.5) as previously described^15,18,19^. For perfusions, mosquitoes were perforated on the posterior abdomen and injected with anticoagulant buffer (∼10μl) into the thorax. Hemolymph samples were placed on multi-test microscopic slides (MP Biomedicals) and observed under a fluorescent microscope (Zeiss Axio Imager).

### Mosquito injections with clodronate liposomes

To determine the effects of phagocyte depletion on the activity of our promoter constructs, 3-5 days old transgenic mosquitoes were intrathoracically injected with control or clodronate liposomes as previously described^19,79^. At 24hrs post-injection, total RNA was isolated from whole mosquitoes and used for gene expression analysis by qPCR.

### Immunostaining of mosquito hemocytes

Hemolymph was perfused from blood- or sugar-fed female mosquitoes at 24hrs or 48hrs post-blood meal and placed on multi-test microscopic slides. Hemocytes were allowed to adhere for 20 min and fixed with 4% PFA for 15 min at room temperature. Samples were blocked with 2% BSA and 0.1% TritonX-100 in 1X PBS at 4°C overnight. The next day, samples were incubated overnight at 4°C with mouse anti-GFP (DHSB-GFP-12A6), diluted by 1:50 in a blocking medium. The following day, cells were washed three times with 1X PBS and incubated with goat anti-mouse 488 diluted by 1:500 in blocking medium for 1hr at room temperature. After five washing steps, samples were mounted with DAPI antifade medium and immediately examined under a fluorescent microscope (Zeiss Axio Imager).

### Flow cytometry

To analyze wild-type and transgenic mosquito hemocyte populations, we performed imaging flow cytometry using the BD FACSDiscover S8 Cell sorter (BD Biosciences). To visualize the proportions of phagocytic immune cells, mosquitoes were injected with red fluorescent carboxylate-modified microspheres (Thermo) at a final concentration of 2% (v/v) and allowed to recover for 30 minutes at 27°C. Hemolymph was perfused from ∼40 individual mosquitoes with an anticoagulant buffer in microcentrifuge tubes kept on ice, as previously described^19^. Samples were centrifuged at 2000g at 4°C for 5min, the supernatant was discarded, and pellets were resuspended in 1ml of 1XPBS. Immune cell nuclei were counterstained with DRAQ5 (1:1,000, BD Biosciences) for 1hr on ice. After incubation, cells were washed once with 1XPBS to remove the excess stain and cell suspensions were transferred to 5ml flow cytometry tubes. Gating was performed using strict threshold parameters as determined by the use of DRAQ5-free (unstained) and bead-free wild-type cells to remove background autofluorescence. For bead-uptake assays, a modified gating strategy was implemented to exclude free beads (based on the signal from a fluorescent bead-only sample) and to distinguish bead signal from the (cell) DRAQ5+ gate. A similar experimental setup was used for the analysis of transgenic mosquitoes injected with fluorescent beads. Flow cytometry analysis of wild-type or transgenic hemocytes with or without fluorescent beads was performed in three independent biological replicates. Data were analyzed with FlowJo v10.10.0 software.

## Supporting information

Supporting Information containing Figures S1 to S12 and Tables S1 to S3

Additional File 1- Promoter Constructs

Additional File 2- Naive mosquitoes IFC

Additional File 3- Phagocytosis assay IFC

## Acknowledgements

The authors would like to thank Emma Howell and David Hall for their assistance with mosquito maintenance, as well as Robert Harrell of the University of Maryland Insect Transformation Facility for his assistance with *An. gambiae* transgenesis. The piggyBac-3xP3-DsRed plasmid was kindly provided by Peter Atkinson. This work was supported by R21AI166857 to RCS from the National Institutes of Health, National Institute of Allergy and Infectious Diseases.

