## Supporting Information containing Figures S1 to S12 and Tables S1 to S3 for "Exploring new dimensions of immune cell biology in *Anopheles gambiae* through genetic immunophenotyping"

### Included supporting information

***Supplemental Figures***

**Figure S1.** Overview of hemocyte promoter constructs.

**Figure S2.** Characterization of piggyBac insertions in transgenic *An. gambiae*.

**Figure S3.** Marker gene expression across different constructs and transgenic lines.

**Figure S4.** Additional characterization of the NimB2-CFP line.

**Figure S5.** Gene marker expression in response to clodronate liposome injection.

**Figure S6.** Visualization of perfused LRIM15- and SCRASP1-GFP cells under native and fixed conditions.

**Figure S7.** Hemocyte-specific marker gene expression responses to blood feeding.

**Figure S8.** Blood-feeding promotes a shift in PPO6$^{low}$ proportions.

**Figure S9.** Determination of gating for flow cytometry analysis.

**Figure S10.** Classification of immune cell subtypes in the P1-P5 DRAQ5 clusters.

**Figure S11.** Determination of gating using fluorescent beads in flow cytometry.

**Figure S12.** Analysis of phagocytosis in immune cells perfused from transgenic mosquitoes.

***Supplemental Tables***

**Table S1.** Primers used for the amplification of the hemocyte promoter regulatory regions.

**Table S2.** Primers used in splinkerette PCR.

**Table S3.** Primers used for gene expression analysis.

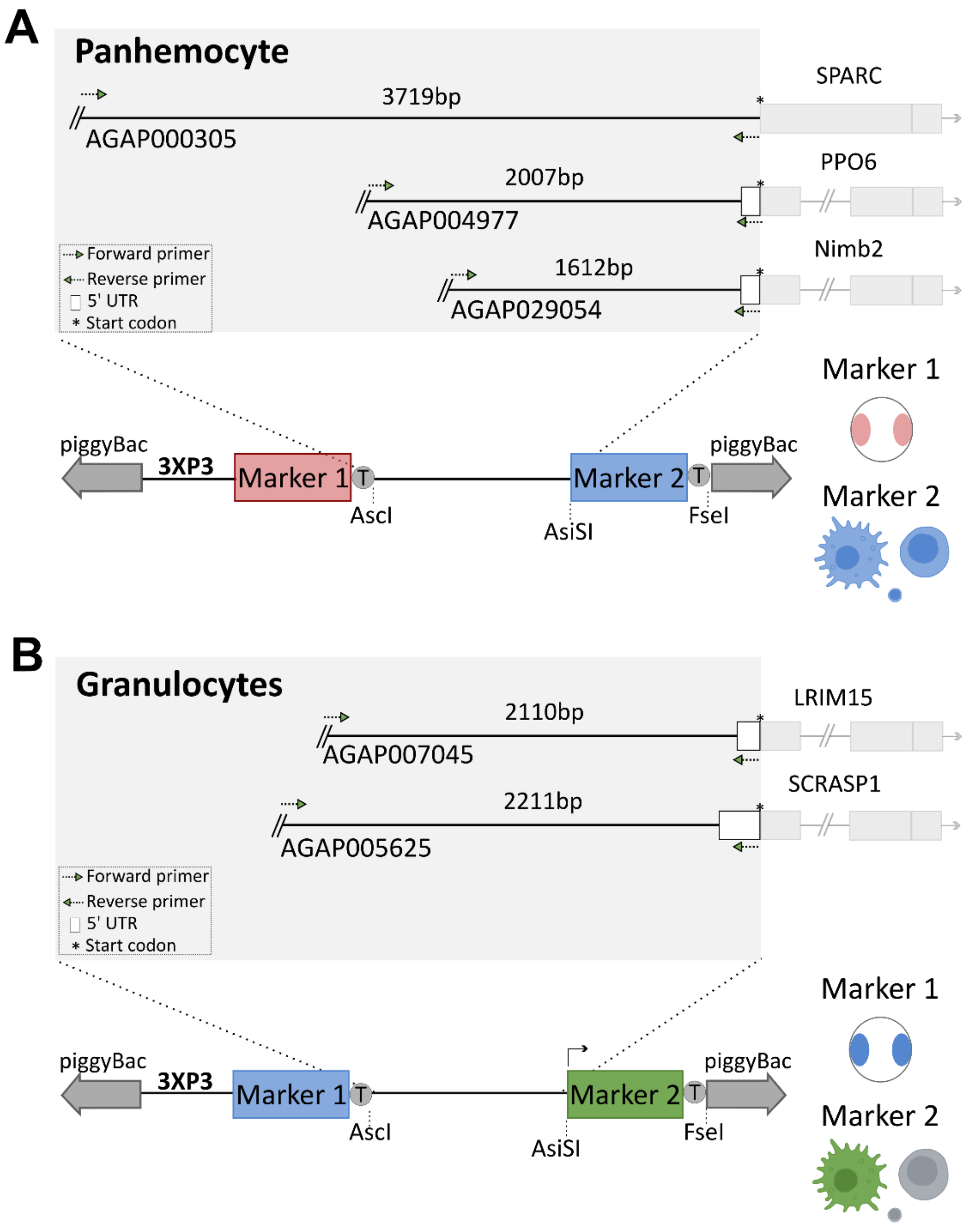


**Figure S1. Overview of hemocyte promoter constructs.** Schematic overview illustrating the constructs used to generate transgenic mosquitoes expressing fluorescent markers under the regulation of promoters that drive expression (**A**) in all hemocyte populations (panhemocyte) or (**B**) specifically in granulocytes.

**A**

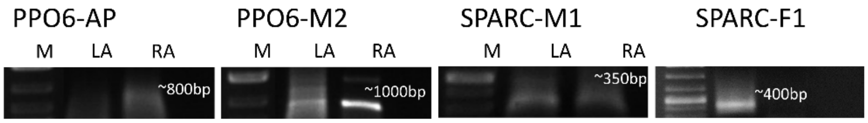


| Strain | Line | Chr | Genomic coordinates | Associated Gene |
|---|---|---|---|---|
| PPO6:CFP | M2 | 2R | 55710203-55710779 | AGAP004403 |
| | AP | X | 9475427-9475283 | AGAP000528 |
| SPARC:CFP | M1 | 3R | 18994307-18994439 | AGAP008826 |
| | F1 | N/A | N/A | N/A |
| Nimb2:CFP | MIB | N/A | N/A | N/A |
| | MIE | N/A | N/A | N/A |

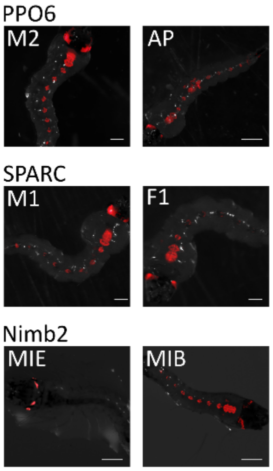


**B**

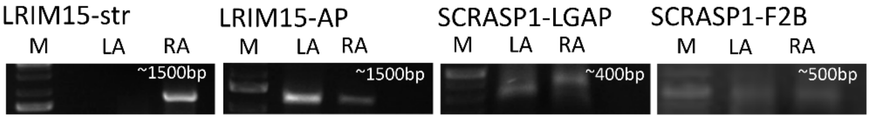


| Strain | Line | Chr | Genomic coordinates | Associated Gene |
|---|---|---|---|---|
| LRIM15:GFP | Str | 2R | 5144656-5146010 | AGAP001446 |
| | AP | X | 11617354-11614600 | Intergenic |
| SCRASP1:GFP | LGAP | 2R | 29680727-29680903 | AGAP002925 |
| | F2B | 2L | 40518618-40518727 | AGAP006994 |

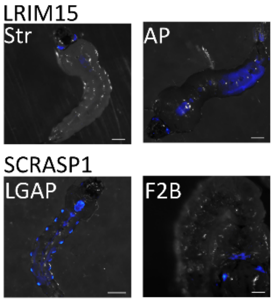


**Figure S2. Characterization of piggyBac insertions in transgenic *An. gambiae*.** Using splinkerette PCR, we characterized the genomic insertions of the piggyBac vectors used for each of the respective hemocyte promoter constructs. Insertion sites and phenotypes produced from the 3xP3 marker (RFP or CFP) are displayed for the putative panhemocyte (**A**) or granulocyte markers (**B**).

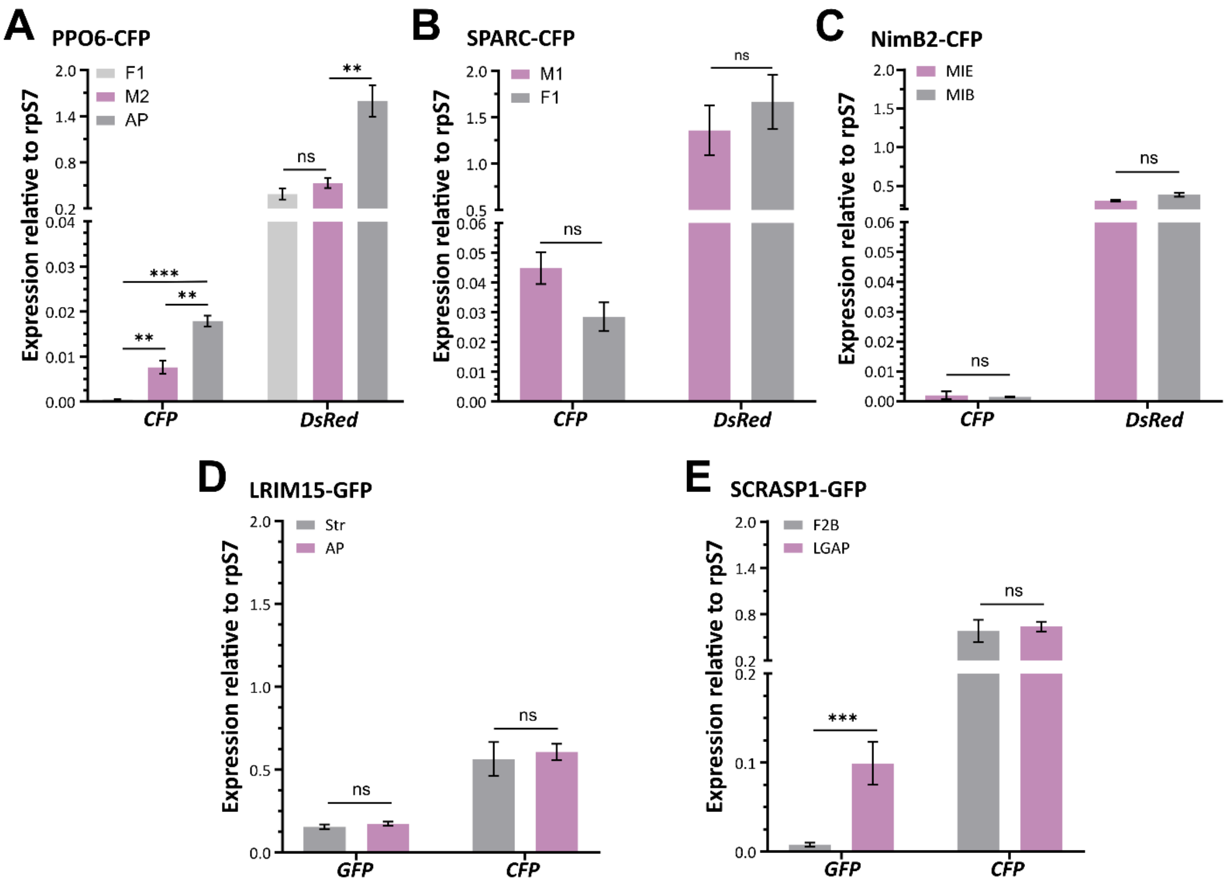


**Figure S3. Marker gene expression across different constructs and transgenic lines.** The expression profiles of each transgene and transgenic construct were examined in 3-5 days old naïve adult female mosquitoes. Panhemocyte marker constructs, PPO6 (**A**), SPARC (**B**), and NimB2 (**C**), were evaluated for *CFP* expression under the control of the respective hemocyte promoter, with expression of *DsRed* to examine differences in the integration marker for each transgenic line. Similar experiments were performed for our granulocyte marker constructs, LRIM15 (**D**) and SCRASP1 (**E**), to evaluated *GFP* expression under the control of the respective hemocyte promoter, with expression of *CFP* to examine differences in the integration marker for each transgenic line. Expression data are displayed relative to *rpS7* expression with bars representing the mean ± SE of three biological replicates. Data were analyzed for significance using an unpaired Students' t-test. Asterisks indicate significance (**P* < 0.05, ***P* < 0.01, ****P* < 0.001). ns, not significant.

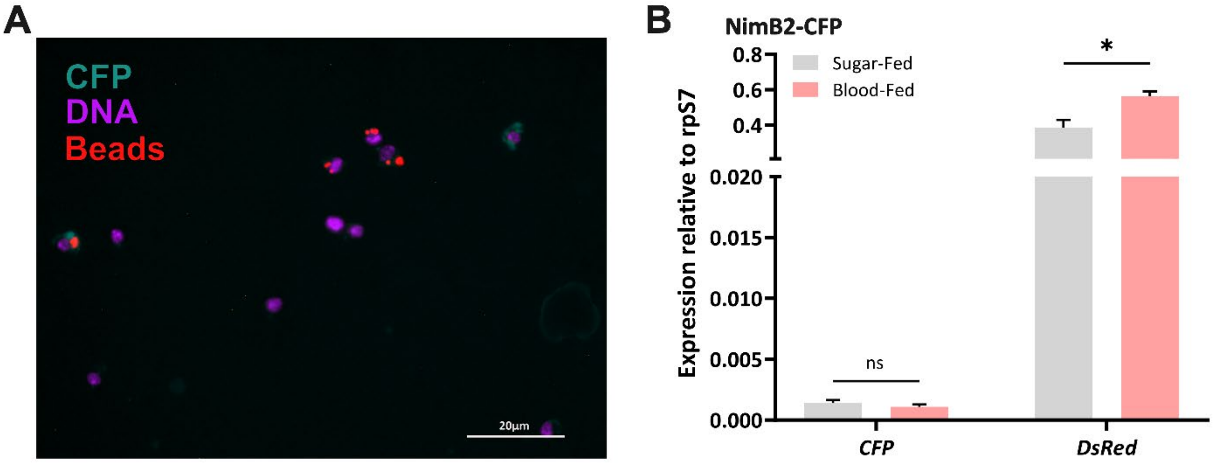


**Figure S4. Additional characterization of the NimB2-CFP line.** The *ex vivo* examination of NimB2-CFP expression in perfused hemocytes was below the limit of detection and unable to distinguish potentially labeled cell populations from background levels of fluorescence (**A**). The injection of fluorescent beads prior to perfusion allows for identification of phagocytic granulocytes. The expression of NimB2-driven *CFP* or of the *DsRed* integration marker were examined under sugar-fed (naive) or at 24hrs post-blood feeding **(B)**. Data represent the mean ± SE fold change expression of three independent biological replicates, with significance determined using an unpaired Students' t-test. Asterisks indicate significance (* $P < 0.05$). ns, not significant.

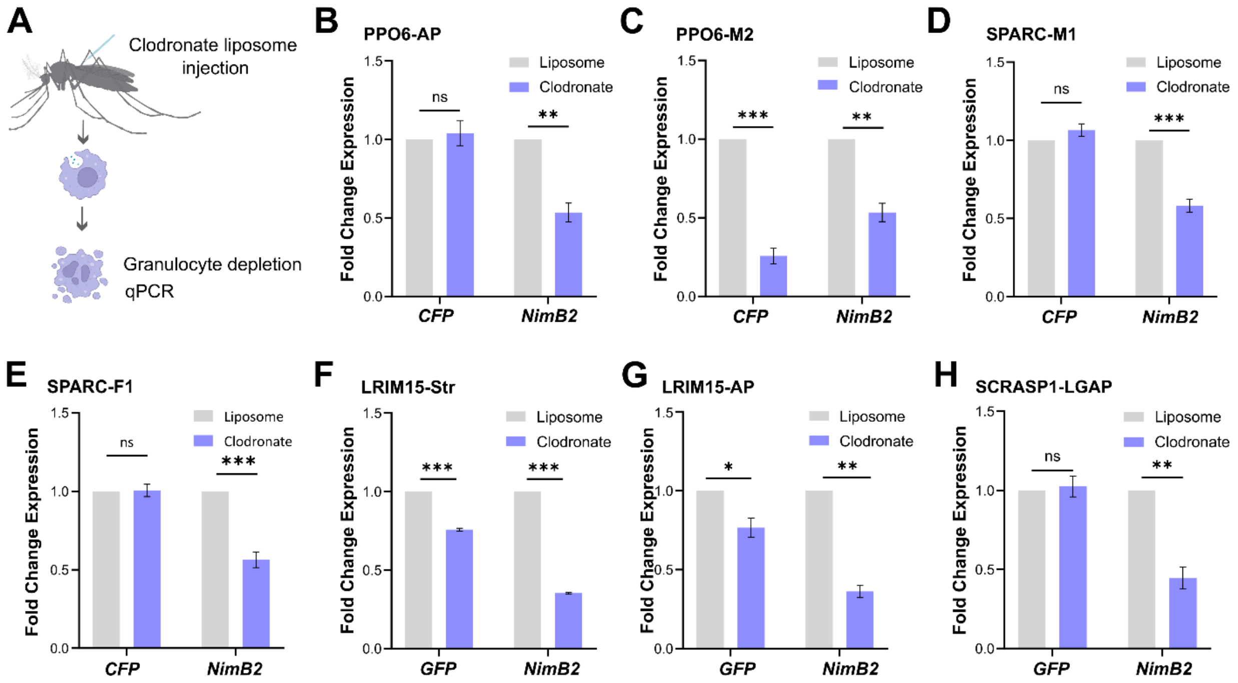


**Figure S5. Gene marker expression in response to clodronate liposome injection.** The expression of gene markers was examined at 24hrs following injections with clodronate or control liposomes to analyze the specificity of promoters in phagocytic populations. The hemocyte marker *Nimb2* was used as a positive control. Data represent the mean ± SE fold change expression of three independent biological replicates, with significance determined using an unpaired Students' t-test. Asterisks indicate significance (* $P < 0.05$, ** $P < 0.01$, *** $P < 0.001$). ns, not significant.

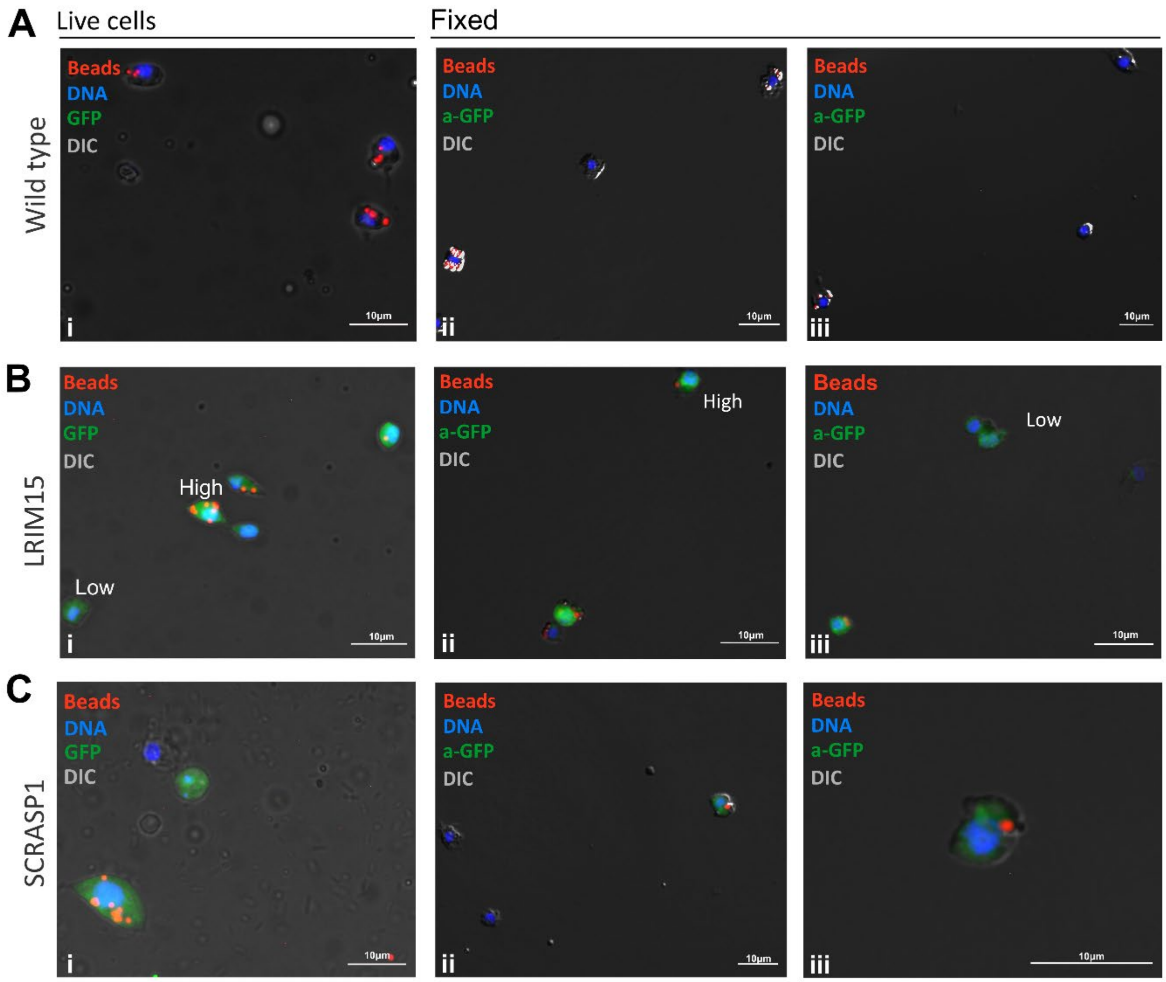


**Figure S6. Visualization of perfused LRIM15- and SCRASP1-GFP cells under native and fixed conditions.** Perfused hemocytes from (**A**) wild type, (**B**) LRIM15, and (**C**) SCRASP1 mosquitoes were immediately observed (i) under a fluorescent microscope or were fixed (ii, iii), then followed by immunostaining with a GFP antibody (a-GFP). Prior to perfusion, mosquitoes were injected with a solution containing 2% fluorescent beads (red) and 1 mM of Hoechst 33342 (blue) to define cell populations and identify phagocytic cell populations. Images were taken using the same exposure time (1000 msec) which was defined based on autofluorescence in the wild type (Keele) background. Scale bar: 10µm.

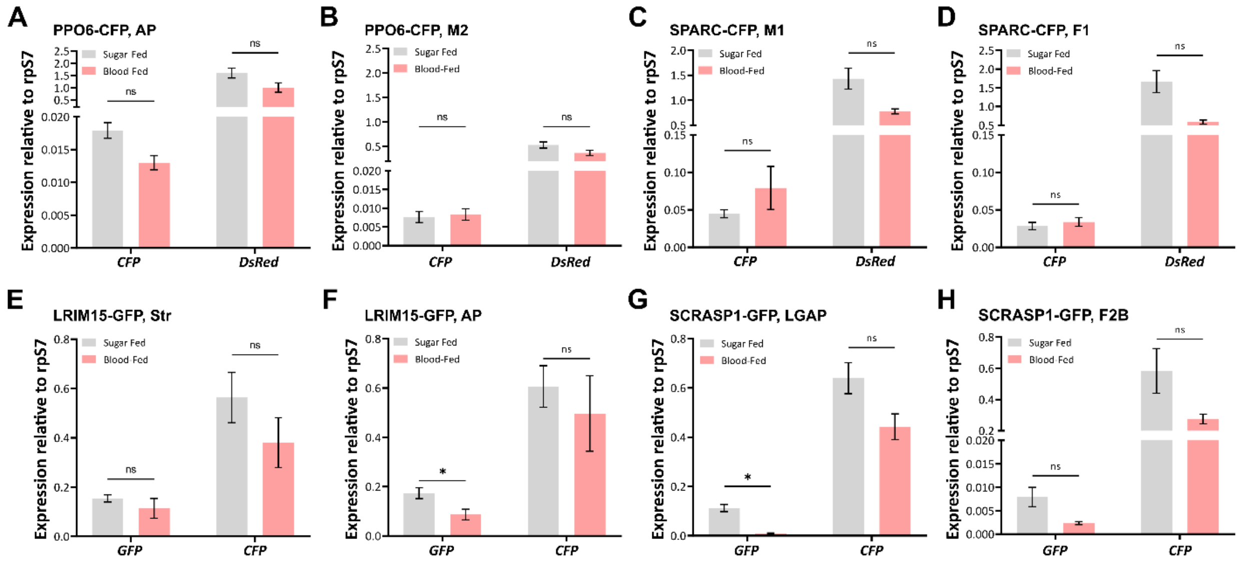


**Figure S7. Hemocyte-specific marker gene expression responses to blood feeding.** The expression of marker genes (*CFP*, **A-D**; or *GFP*, **E-H**) were examined for each of the hemocyte promoter constructs under sugar-fed (naive) or at 24hrs post-blood feeding. In addition, the expression of the integration marker (*DsRed,* **A-D**; or *CFP*, **E-H**) was used as an internal control. Data are displayed as the individual transgenic lines for each transgene construct for PPO6 (**A-B**), SPARC (**C-D**), LRIM15 (**E-F**), SCRASP1 (**G-H**). Expression data are displayed relative to *rpS7* expression with bars representing the mean ± SE of three biological replicates. Data were analyzed for significance using an unpaired Students' t-test. Asterisks indicate significance (**P* < 0.05). ns, not significant.

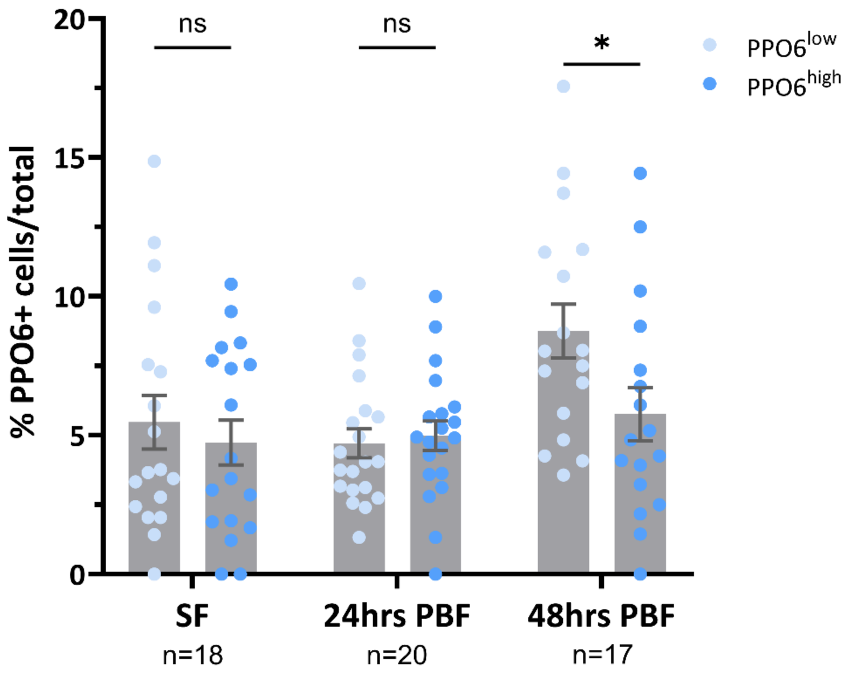


**Figure S8. Blood-feeding promotes a shift in PPO6$^{low}$ proportions.** The percentage of PPO6-CFP cells with low (PPO6$^{low)}$ or high (PPO6$^{high}$) CFP fluorescence are displayed as the percentage of total hemocytes. Data from individual mosquitoes are represented by dots and represented as the mean ± SE of three independent biological replicates. Statistical significance was determined by a two-way ANOVA followed by Sidak's multiple comparison test. Asterisks indicate significance (*$P$ < 0.05). ns, not significant; n numbers of individual mosquitoes examined.

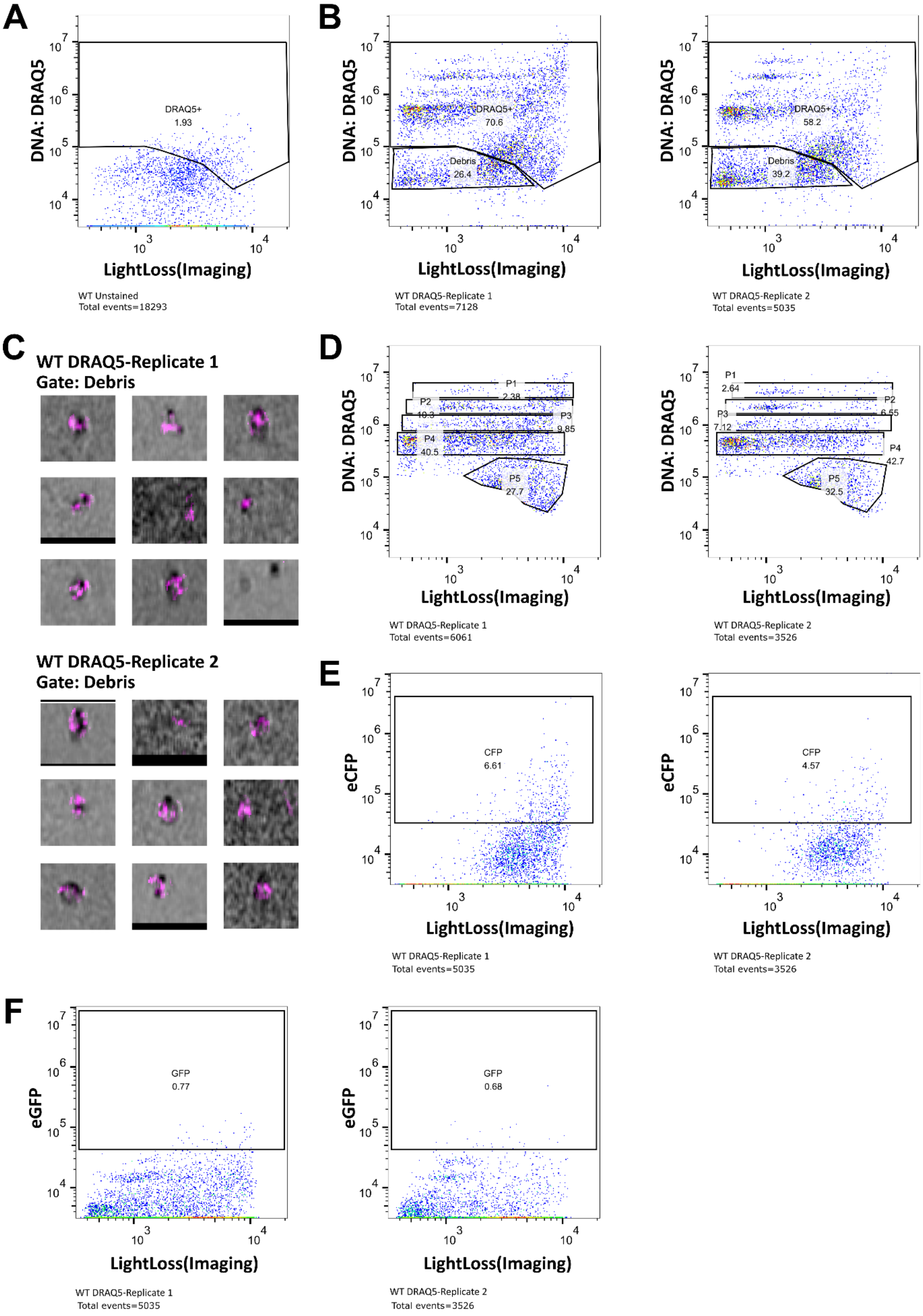
A
DNA: DRAQ5
LightLoss(Imaging)
DRAQ5+
1.93
WT Unstained
Total events=18293
B
DRAQ5+
70.6
Debris
26.4
WT DRAQ5-Replicate 1
Total events=7128
DRAQ5+
58.2
Debris
39.2
WT DRAQ5-Replicate 2
Total events=5035
C
WT DRAQ5-Replicate 1
Gate: Debris
WT DRAQ5-Replicate 2
Gate: Debris
D
P1
2.38
P2
10.3
P3
9.85
P4
40.5
P5
27.7
WT DRAQ5-Replicate 1
Total events=6061
P1
2.64
P2
6.55
P3
7.12
P4
42.7
P5
32.5
WT DRAQ5-Replicate 2
Total events=3526
E
eCFP
CFP
6.61
WT DRAQ5-Replicate 1
Total events=5035
CFP
4.57
WT DRAQ5-Replicate 2
Total events=3526
F
eGFP
GFP
0.77
WT DRAQ5-Replicate 1
Total events=5035
GFP
0.68
WT DRAQ5-Replicate 2
Total events=3526

**Figure S9. Determination of gating for flow cytometry analysis.** Prior to flow cytometry analysis, controls were performed to apply the proper gating of cells for further analysis. Unstained cells from wild-type *An. gambiae* were used to set threshold values (**A**) for gating for DRAQ5$^+$ positive events (cells) (**B**) to remove any autofluorescence background or cellular debris (**C**). Wild-type cells exhibited five distinct DRAQ5 signal patterns (**D**), corresponding to subpopulations P1-P5 that differ in their DNA content. Additional controls using wild-type DRAQ5$^+$ cells were used to set threshold values for gating CFP$^+$ (**E**) and GFP$^+$ cells (**F**). Of note, wild type cells contain a relatively high levels of CFP background fluorescence. For each experimental condition (**B**-**E**), data are displayed for two independent biological replicates.

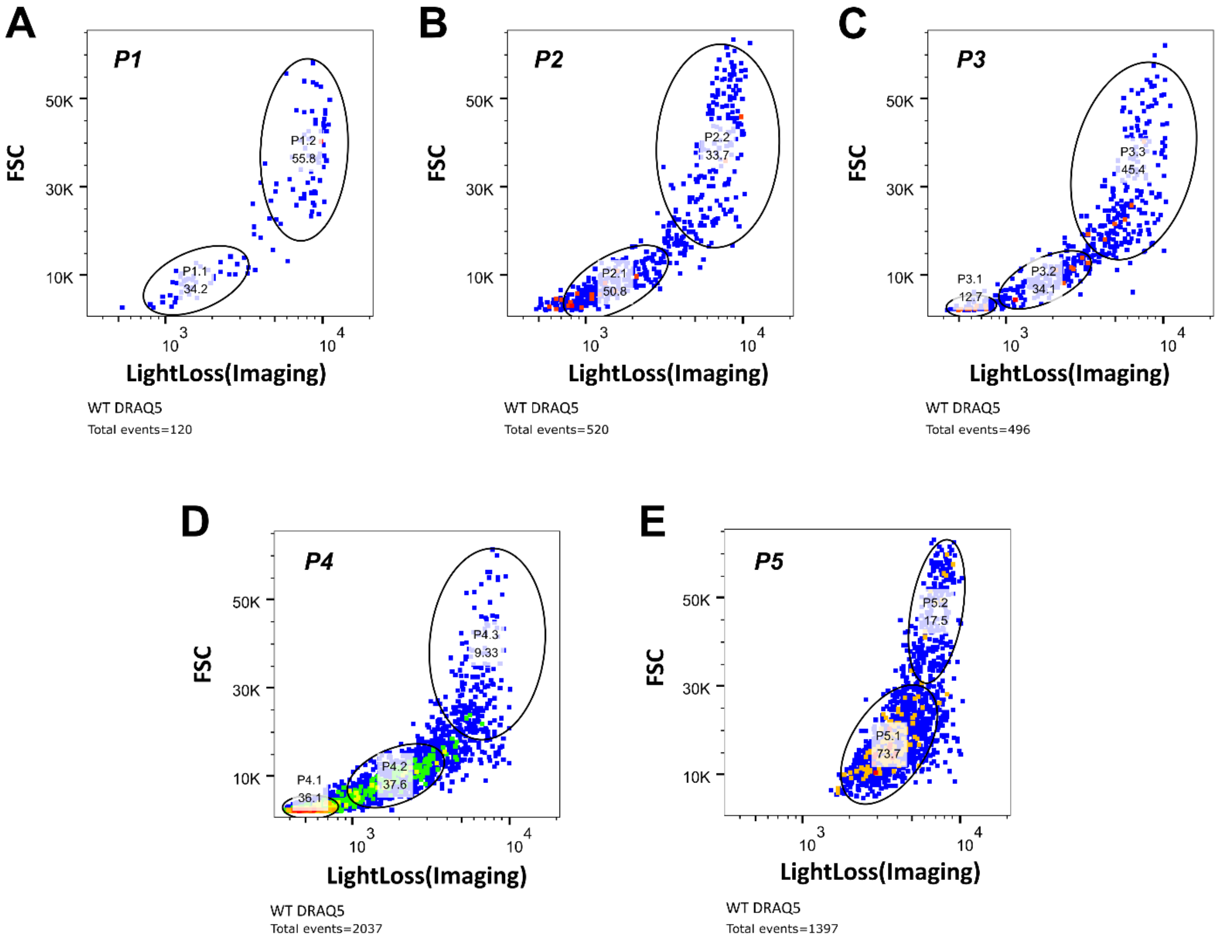


**Figure S10. Classification of immune cell subtypes in the P1-P5 DRAQ5 clusters.** Initial sorting of DRAQ5+ cells identified 5 cell clusters, P1 through P5, based on DNA content. A further examination of the P1 (A), P2 (B), P3 (C), P4 (D), and P5 (E) clusters by size (FSC) and axial light loss reveals additional subpopulations for each cell cluster. Subpopulations are defined by circles, with the percentage of cells displayed for each subpopulation (of total).

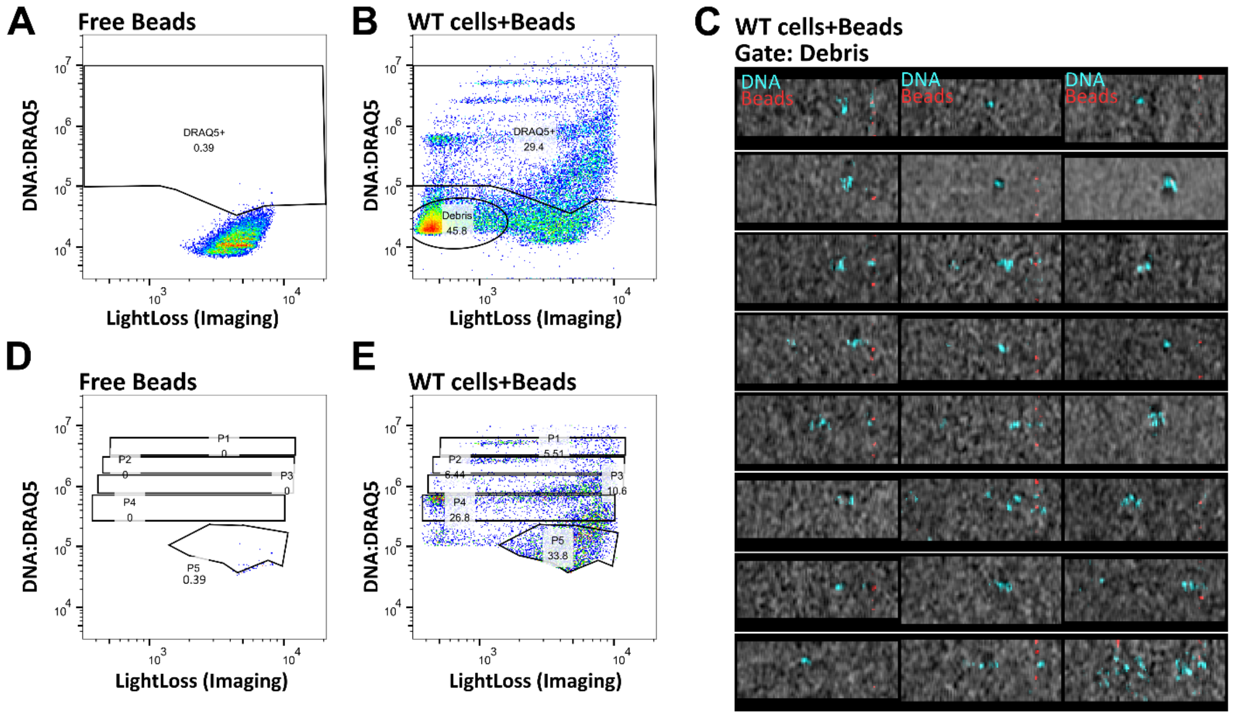


**Figure S11. Determination of gating using fluorescent beads in flow cytometry.** Prior to flow cytometry analysis, controls were performed to apply the proper gating of fluorescent beads for further analysis to examine phagocytosis. (**A**) A fluorescent bead-only sample was used to distinguish cutoffs for bead (red) and DRAQ5 (far-red) signals. (**B**) Mosquito perfusates were examined after the injection of beads to identify phagocytic cells. Cells were stained with DRAQ5, with events identified as immune cells (with or without beads), free beads, or cellular debris which was confirmed by imaging (**C**). Using this gating methodology, no bead signal was detected within the P1-P5 groups (**D**), while enabling the gating of immune cells (with or without beads) according to DRAQ5 signal (**E**).

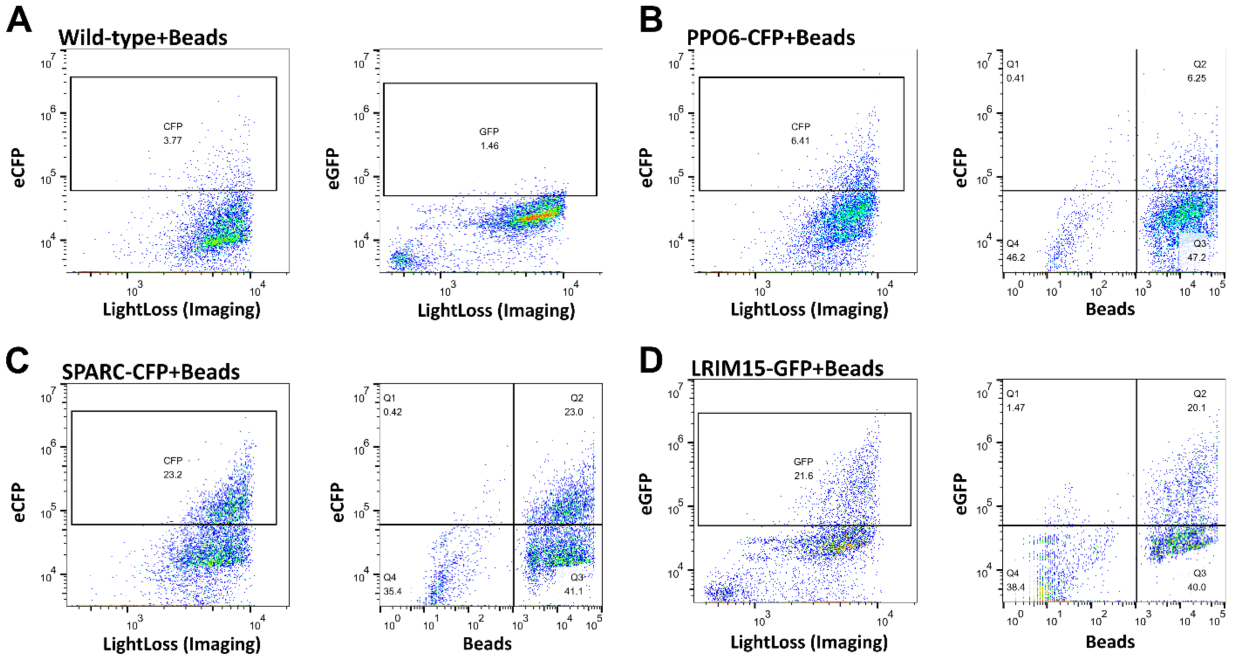


**Figure S12. Gating thresholds for analysis of phagocytosis in transgenic lines.** Prior to flow cytometry analysis, controls were performed to apply the proper gating of CFP and GFP fluorescence to perform phagocytosis assays in PPO6, SPARC, and LRIM15 transgenic lines. (**A**) Gating thresholds were set for CFP and GFP fluorescence in wild-type mosquitoes injected with fluorescent beads. When these gating strategies are applied, phagocytosis was examined in PPO6-CFP (**B**), SPARC-CFP (**C**), and LRIM15-GFP (**D**) lines. Phagocytosis assays were performed in three independent biological experiments with data summarized in **Additional File 3**.

**Table S1. Primers used for the amplification of the hemocyte promoter regulatory regions.**

| Promoter | Gene ID | Forward (5'-3') | Reverse (5'-3') |
|---|---|---|---|
| SPARC | AGAP000305 | gtacggcgcgccGCAATCACATCAGCTTCAAGAAG | gtacggccggccactagtgcgatcgcCGTTCGTCGGCCCGTTTC |
| PPO6 | AGAP004977 | gtacggcgcgccTCATCGCTGGGAAGAATAAAGGAAG | gtacggccggccactagtgcgatcgcTTGATTATCCGTTAGTTTTTCCACTTAAAG |
| LRIM15 | AGAP007045 | gtacggcgcgccAATTGGGAGAAAACAACAGAAGAGC | gtacggccggccactagtgcgatcgcTTTTCAAGGGAAACACTATGGACAG |
| SCRASP1 | AGAP005625 | gtacggcgcgccAGATGCACGGGGATTAAAACAATTC | gtacggccggccactagtgcgatcgcTTTCGCGCAAATCTTCGCTTG |

Small letters correspond to restriction sites attached to the 5' end of each primer following the GTAC spacer sequence to enable the digestion of PCR products.

**Table S2. Primers used in splinkerette PCR.**

| Primer | Primer sequence (5'-3') |
|---|---|
| Splink-GATC-Top (BglII) | GATCCCACTAGTGTCGACACCAGTCTCTAATTTTTTTTTTCAAAAAAA |
| Splink-CGG (MspI) | CGGCCACTAGTGTCGACACCAGTCTCTAATTTTTTTTTTCAAAAAAA |
| Splink-Bottom-Universal | CGAAGAGTAACCGTTGCTAGGAGAGACCGTGGCTGAATGAGACTGGTGTCGACACTAGTGG |
| piggyBac LE#1 | CAGTGACACTTACCGCATTGACAAGC |
| piggyBac LE#2 | GCGACTGAGATGTCCTAAATGCAC |
| piggyBac RE#1 | CGATATACAGACCGATAAAACACATGCGTC |
| piggyBac RE#2 | ACGCATGATTATCTTTAACGTACGTCAC |

**Table S3. Primers used for gene expression analysis.**

| Gene | Gene ID | Forward (5'-3') | Reverse (5'-3') |
|---|---|---|---|
| CFP | AGAP000305 | ATCAGCCACAACGTCTATATCACC | TGTGGCGGATCTTGAAGTTGG |
| GFP | AGAP004977 | AACAGCCACAACGTCTATATCATG | TGTGGCGGATCTTGAAGTTCA |
| DsRed | AGAP007045 | CGACATCCCCGACTACAAGAAG | GTAGATGAAGCAGCCGTCCTG |
| rpS7 | AGAP010592 | ACCACCATCGAACACAAAGTTGACACT | CTCCGATCTTTCACATTCCAGTAGCAC |
| NimB2 | AGAP029054 | CAATCTGCTCAAATGGCTGCTTCCACG | GCTGCAAACATTCGGTCCAGTGCATTC |
