## Additional File 1- Promoter Constructs for "Exploring new dimensions of immune cell biology in *Anopheles gambiae* through genetic immunophenotyping"

**Extended File S1: Promoter sequences**

***List of putative hemocyte-specific promoters***

Putative promoter sequences of genes-markers expressed in *Anopheles gambiae* hemocytes. Letters highlighted with gray indicate the 5’ Untranslated Region of each gene.

1. **>AgNimb2**

ACCTCCTCCACTACCACTTCAACAACCGAAGCTACTGGACAAATGCTACCAAACGCGACCTACGCCGAATCGTACCACGAGTACCTTCCCAACAACACCGAGATGAGCATGGTGCTAACAAAAGGTCACTCACGAACTGCCAGCAACGAAAGCATGTACGAGCTGGTGACGACTACGTCACGCGTCGAGATCACCTTCATCGATACGGCGGTGAAGAAAATTGAGGATCCCTCCACGATCAACCATACAACGCCGGAGGAGGAGCCCGCTTCCCTGTCTGTAGAAAGCTCCAACGAAACGGACACCCAACTGGTGTACTTGGAATCGGAGAACCATCCCGTTCTGGTGGGCATAACCGAAGATGGGAGCGTGCAGAGTGAGGCAAATGATCGACAGGCGTTCGTCACGATCTCCTGCTTGCTGATCACACTCACCCTGCTAATGTCGGTGGTCATCTATATGAAGCGTTTGAATCGCAAAACGCTTGCCAAACCTGCCGCTAGTGTGCCACCGCCGGTGAAGATTGTTGACGAGAGCGTCCAAACGGGGAATGATTCGAACCGCTCGTCGGATCCACTTCCCGATCTGCCCCATGTCACCTACACGCGGGTAAAGCCAAAACTGCAACGAAACGGTGATATCGGTAAGTGTATATAGAAGCAAGTTCTCCTACAATCTAACAGCAAGGCGAATCGTTAAAACTAACTATTATCTTCCCAGAACACTATGATGTTCCGCCGAATAATAGTTTCATCCATCGTGCAAAGTCAGCATCCCCTTACAACTACAACTTCTCGGTCAACCAAAAGCAAGCTCCCAGAAAGTATAGCCTTGAGCATATCTACGACGAAATTCAGTATCCCCCGTTGGCGGAACTGACGAGCAGTACAGCAACGCAGTCCGATAAGCCGACGTTGCAGCCGGGTGAGAAGGAAGACAGTTATTCCAAACCGATGATTCTGGCGTAGGATAGGGCTAAGTTTGCTTATATGTGGTTCTTATGTTGAACTAAATAAATGTATTTTTTTAACTTACCATAAAATAGGAATTGTTGGTTACAGCGTGAGTATCGTCCATCTTCAATAATAGCTACCTCGTCTAGAGTTTAAATTATCTCAATTTTTTTTCTCTTGGCATTCGCTACGGCATGTATTTCTCACTCTTCCTTGTGGCCGTTGCGCCACAGGTGACATGAAATATTAATTTCTATATCATCCATCAATCAATCGCTTCACAGCGCGCAACTTAATACATTCCATTCAGGATTCCCCTTCCATGCCTTTGCATTAAACATCTCGCATCACAATACATCAAACGTGAGTATGCCTGATGGGGCGCAGATGCCCGGTGGCTTACTGTGTTGCGCCACTTCAAATCGCACCCGAGATCGGACTTTTAAGAATAAGCACATCACGTCATACGAGACGTTCGCCCACACACCACAACGTTTCGGTGCGATATCACATCATGCCAAAGAACGTCACCAGCCGTACCGCGACGAGTTCTCGCCATTAGGTTCTACTGACCGTTTACTTTAATCCATCGCGAACCTTCCGTGGGGTAAGATTGAAGTTGAAGCTGTAGTGACCCCATTTCTTTACGATATTACACC

2. **>AgSPARC**

CCGCAATCACATCAGCTTCAAGAAGTACACGAGCCCCCCCTTCCTACATTAACGACCTTAGCAGTACTTTTTGCGTATCAAACAAACACACACACACACTAACACATACAGATCGCCCTTAATTACTTCACCGCATGTGCAAATGTCGGCAAGCGAATGAGGTTTGCAGGGACGGCCGTCCATCCGGTCCATTCGAAACTTCAGAACACACTTCGGCACATTCTTCGGCTGAGGAAACGAGCGCGGGAGCGGGCGCGCGCGCACGCAATCTCACATCTCATACACGAGCTAATCATCCGGCGGGGTCTTATGGTTTTGTCTTTAACAGTTCCCGCCCGCCCTTCCCCAGACCCCGCCGACTGTGGAGGGAGTGTGGCGAGTGTGTTCTCGCTTGCTCCGCTTGGCCGGGAAGAGTCGTCGCATCACCGGGACGCCCACGAGATCCCATCCCGGCGGGTTCGGGTCGACTCTCTGTGCCCGAGTCGAGTCAGTCTGGCTCTAACGATCGTGCCGATCGGACGCCACCAAATCGGGTGCAAGAGACGTGCTTCCGGAGCTACCGCAGCTACCGTGTAGTGTTATTGTTTACTGGTGCACCAAGTCCTTGGGTCTTTGGGAACATTCTGTGCTTCAACGATATTCACCAACAAGAAACAGAAATCTTCAAAGTAAGTGTAGATTAAAGTTTGGAGTAGCTGGGTGAGAGATGTTGGGAACTGTGACCTTACTCCTCAAGCCTGTTGGGAGTCGTTAGTCCCCTGGAACCCTATTCAAAGTAAGTTCTGTGTACTGGGTTTGAATTGTTCCACGCTGTGCCTAGCAATGGTTACATCTTCAATCGGACCTTATGTGGTCGCTTATCGCGGTCCGGTACGGATGCGACAAGTTCCTTCCAACGGTCCAAGCCTAACCTTCAATCAAACTGATAACCTCGCGCGCTCTGAAACCGCGGCATCTTGCGGCGGAAGCAGTCTCCACACCGAAGCTGCCAGTGTTACCGAATGTCGCCCCGAAACGGTACACAGGATGTTACTTCACACCACAGACCAACGTCAGCAATTAATCGCTTGCGCAAACGTCTCCTCTGCCAGCGCCAACCAGACACCCTCACGGGGGATTTTGAAACTGTAAACGCTCCCGGACGGTTGTGGCAGGACAATGTACCGAGGGGGGAGAGGGGGGGGGGAGTGTACGGATACACCGATGCTCAACTCGGTACAGCACTGTTTACAGACAACTCGGGTGTCGTTATAAATAACCCACCGGTGGAGGTTTCGGTTGGTTTTCGGTTTTTGGAAACCTCTCCATTGCGGGTCCGATACCGAGAATCAAAGGGTGATGTCCGGGTAAGGGGACGGGATTCCGAAGGCAGGATCTCAAGTGCATCTGCCGACGAGGGGACGGGCTACGTTAAGCTATGTCTATGCTTCACTTTAACGACCACTCATTTCCCTAAGTGCAGCGTGCGGTCGGCAACTCCAGAAACTCCATGCAAAGCCCTGCTAGGGATTGTTTGGATAATATCGCTCATCAAGCTGGAGAGTTCGAGGGCGGAGATTCATCATCAAAACAGTACTTCTGGTACAGTATTATTACTGTTTGTCAGTGCAGTGGTATGTCCCCTTTCGGTTCCAGGGGGTTTGTCCCTTTCAGCGAACTCTCAAACCCAAACAATCTCTAACGAAAGCATAACCGAAAGCAACCGCAACAAAACACAAACACTTGCCTCTCCGTTGTTGCCACGCGCTTCTCCTAAACGCATCGTATGACATCAGCATTTGGCAGGACGGTTGTCGGTTTGATTTCCTTGCCCATGTGGCCCGTTTTGCATCTTGCGCTTTCCTGCGCTCCGCACTTGCTACGTCGCAACAAAATTCGCTCAGACTGTGTTCGCTGTGTGTGTGTGTTTTTTATCGCATCGGAAGTACGCTGCGGTTGCGCCACTTCCGCCTAGAACGCCCTCCCCCACTCGTTCTGACACAGATTTGTGGCACATGTGTGCCTTGGCTAGAGCGGCGACAACGTTGTAGCTTTAATAAGCAGCTGGCTGGTGAGATTTTGCTTCCCACTTCCCACTCCATCGCTCCAATGATGATGATCTCAGAATGTTGCTTTTTTGTTTGCGGTGCCTTTCTTTCTTCTTGTTGCTCCAGTGCGCTTCGTAACTGTCGCTCCAATAATATGCGAGCCTGGACGAATGAAGCGTGCGGGTCTTTGCCGGTCCCGGCGCAGTGCACAAACTGCCAACATATGCACTGGCGAAACATTGTTTCTTGCCCGTGGTCATGCGGAGCCGTGTCAGCGTTTTATTACCCGTCCGGGCAGAGATTGCATCATCGCGCAAGGAGGGTCTACCATCTCGGTCTACCATCGACCGACCGAGTGAGCAAGTTTTACGATCTTAAACCAATCTTAATTACCGCGGCGAGCGCAACGTGTTAGGGCGGCTTCCTGATACCGCCGTTTGCAATTCGTAGCAACGTAAAAGATGATGCACCATGGTCCTAAAAACGCATCTGTTTCCTCGCACACACATGGAGCCTTGTGACGACTGGCACGTTTTAATGATGCAGAACGAGTGCTGAAAAGAAAGGTCGCCAACTGGAAGATAGGATTGAAAGGATTTTTTTTTTAAAGAGCAAACTGTTACCTAAAGTGTGTCAGATACGGGTTAAAACGTGGAACCTTGGCAGATAATTGAACTCTTCAAACTTCCGCAAAGTCCTACAGGCACTCAGTGTCCCATCATTCGCATTCTTGAAGATCTTATTGTCGTACATCAATTAGGTATCTAAAACAAATACCTTTGGAGTCGCTCCCTAGACCTTAGATCTATCTCTTCTGACTCGTGCCTAAATAACTACGTTCTTATCTTATCCAAGTACTTAGACAGGAACGTTTGGACAGTTCCTTTAGCCTCTCCCAACCCATACTCTACTCTCAGACTTTAACTTCAAATTCTCCACCAGGACCTAAAAACTGGCCTATAAAGATCAATGTCTAGCAAGATTGTATCGCGAAATTGCGCAAATAATTTATTTTTGTTGTTGCAAATAAATCCGTTGCATTTTATGAACTCTGGAAGAGATTATTATCGTCAAACGATATTAGCATCTCATGGTCGCTGTACTGATGGTAGTACTTGGTACAATTACTTGGCTTAAGTTGTACTCTCTCTCTCTCTCTCTATCTCTCTCTCTCTCTCTCTCTCCTTCTCTCTCTCTTTCTGTCTTTCTCTCTCTGTCTCTCTCTCTGTCTCTCTCTCCTTCTCTCTCTCTCTTTCTCTCTCTCTCTTTCTCTCCCTCTGTCTCTCTCTCCTTCTCTCTTTCTCTTTCTCTCTCTGTCTCTCTGTCTCTCTCTCTCTTTCTCTCTCTCGCTCTCTCTTTCTCTCTCCCTCTCTTTCCCCATATCTCTCTCTTCCTCTCTCTCCTTCTCTCTTTTCCTCTCTTTTCTTCTCTTTCTTTCTCTCTCGTTCTCTCTTTTTCATTCAACTTCTCTCTCTCTCTCTGCCGCTCTATCTCTAATTGTAAATGCAATCAATTGTAAATGTCAACGCTATTTCGGTCGAACCTTATCTTGGGGTTTAGCCATTGGTCCTTCGAACCGGATTTTCCACTGGCATTGGCTCTACTAACCATCTAACTTCACCTTCTCTCACCTTTCTTTCCCCCTCGCCGGCGTCTTGCCCCAGCTTTTGTGAAACGGGCCGACGAACGGCG

3. **>AgPPO6**

CCTCATCGCTGGGAAGAATAAAGGAAGCCATTCAGTCCGGTTTCGCAATGGCAGTATGTATTGTTGTGATATAAAAGAAAAATTTTACCTTTTGGCAAAACCAAGCTAATGTTTATTACAGGCGGACGGGACACGTGTTCCTCTGGATCCTAAGAAAGGCATCGATATTCTTGGCAATATTATGGAAAACTCGATCCTTTCGGTCAACGTACCGTACTACGGTAATTACCACTCGCTTGGCCACGTTCTCATCGGCTTTATCCACGATCCGGACAACCTGTACCTCGAGGGACACGGTGTGATGGGTGACTTTACGACGGCAATGCGTGATCCAACGTTTTACCGTTTCCATGGCCACGTGGACGATGTGTTTGATATGCACAAGCAAAAGCTTTCGCCATACAAAGCGCACGAACTGTCCTTCCCAGGTGTATCCATCTCGGACGCAACGGTGCAGATTACGAGCGGTAAGGCGGCCAGAAATAGATTGCTAACCTTCTGGCAACGGACGCAAGTTGATCTGGGAACGGGGCTAGATTTCGGACCGCAAGGTAACGTGTTGGCAACCTTCACCCACATCCAGCACGCACCGTTTGCGTACCAAATTATGGTACAAAACGAAACGGCGGAGCAAAAGAAGGGAACTGTTCGCATTTTCCTCGCCCCGATCTACGATGCGAACGGAGAGCAACTGTTACTGAGCCAGCAGCGTCGGTACATGCTGGAGATGGACAAATTTGTCGTCAAGTGTAAGTATACATTAAGTTGAGCAATACTGTTATGGTGCTGCAATCGTATCGTGTGTTTCCTTCAAGTACATCCTGGCGATAATCGGATCATTCGACGATCGGACCAGTCAAGCGTAACCATACCGTACGAAAGGACCTTCCGGCGAGTTGACGCTTCCAACATGCCGGGCACGGAGAGCTTCCGCTTCTGCAACTGTGGCTGGCCCGATCATATGCTGCTGCCCAAGGGACATCCCGATGGTCAACCGTTCGATCTGTTTATCATGATTTCTGATTACAAGGACGATGCTGTAAGCACCGGATTCAATGAGTGAGTACATTACGGAATCAGTATTGATGAACTCGGTTTTAACTTTTGAGTTTTTTTTTAAATATCTTATGCTCATGTAGGAATGAAAACTGTAACGATTCACATTCATACTGTGGTCTACGCGATCAGCTGTATCCGGACCGTCGTGCGATGGGTTTTCCCTTTGACCGACAGCCGGTTGCCCAGGATCACTTGATGAAGGACTTTGTGGGCAGGTTCCCCAATATGAGTCGTACCGTAGCGGAAGTTATGTTCACCAACACTATCATTTCACGCACGTAAATGGCATCACGATAACCGATCGATGAACGATACTTGTGGCTGACCCGTATGATCGTTTTTTTTTATCAACGAGGAACATTATATTAAAATGCAATGGAAAAATGAGCAATAAAATATATTCATAATCAAGCAAATATATACCGCATTTCATTGCATTCTTAAAGTTATTTCTATTTTTTGCAAGTATTATGATTTTTTTCAACAATGCGTGTTATTTTCCTCAACCACCGATATGCAATATGGATAAGGCACACACATTCCCAGTTATCGTAATCGCTTACCAAATGATTGGAATACCGCAAACATCCACAAAAATAGAACGGATTGATATGTTTTATCAAAGCTTCATGCATGTGCATACGCGTGGTTTACATTTGCTTCCCATGTACTGCAACAATTGCAGATTACAATTGCATTGTACCATATCATTATCGCAGATGCAGTGGTCGTAAACCGCAAAAATGGCGATTGGAATTGGAAGAAATTGCATACAACCAGGCACCGGCACGTCAAGTGCAAATCAGAATGATATATAAACTATACCAACTACTTCTGTAGCATCACAAACGTGACTGTAACAGTGACTGGTGGTTCTCGTTTCGTTCTCGCGCGTCTTTAAACACCGAATCGTTCCTTTAAGTGGAAAAACTAACGGATAATCAAGCG

4. **>AgLRIM15**

AATTGGGAGAAACAACAGAAGAGCGCCTTACGTTTGTGCCTTAAACTGCTTGTTGTGGTTTGATACGAAAAAAGGAGAGGAAAATGAAAATGTGTGATAAAATGCGTTTTCCCATACGCTTTTCCTTTCAGCAGTAGTAGCTTCTATGATCGTTGTGATAACATAAGAAAAAAGAGACGTATTTTAAGAGCCAATGTGGAACCAACTGCGATAAGCATACCTACGATCGCGTTGAAAGAACAACAGTTCAAAAGGTGGTGTGATGAAGCTTCACGCATTGAAGGTTATCGTGTGTTTAATTTGAAAGCGAAACATCTTTATTCTTCCTTTCAATTCCTTTCATTCACGGTATTGTCTATCTTGCAATTGATTTGGCTTAGGCAACCAAAAAACATAATCGATGATCGAACGCTTGATAAACCATATAAAGAAGAGTTTGGAGTGCAAAATGTCGAATTGATGCACATGAATGGATTTGCATGAGCATAGATTGCAGCAAACTACACATAATTAATCTTAAGGCTGTATAGAAATAAAACTTATTCACATAACACATAATTCACATAATTTTCTCTCAGATATTTGCAACTTGTAGTTTGACTTAAGCTACACATTTCTAATCAATTTTACACAACAACTCAATTCATTGTAATCATTTTATTTCAGACATTTGGTTAAAGCTTCATAATGTATACCAAAGAATCTTTTTCATGTAATTCGTTAACATTATACGAAAATTACATTATACTATAAGAGTCATACACATGTGTTTGAGTTTTCTAAAGGTGAGAAGTTTTTGGGGCAAGTAAAATGCATTGCCGGCATCGCATCATGTTTGCTATACCAAAGCTTTGAATAACACAAAATTATTCGAAAACCGTCTGACATAAAAAAGAGAAAGAAAGAAAACTCATAAATGCTCCAATAAGCATCCAATTAATAACGTGATGTGAAATGTAAAAGAACCACCCATATGTTAAGGTGACCAACAATTTAGAACGGTCTAATCCGCGAATACATGGCGAACCCGAGAAAGCTAAATTAAAGTTTCATATAGTCAGAATTTTGAAGATTACAAGGATTTCCCACGAATTTGACACCAACACAGATCCATAAGACATGTAGGTGCACACCCAATACTTAGTATGCTATTCGAATTTTTGTTAATCAAAATCAAATCACATCCAACTGTAAAAAGAGAACATTATCATTTAAAACTATGGACTTTTCACATTGGATGCACACTGAGCGTTCCTACCCAAGGTAATGATTGATAGCTCCAACGTTTATTGTGACCAATCTGTTCATGGACATCCAAGATCGACTGACCTATCACATTGCCAACAGATCTAAGAAAAAATCGCAATTTTGCACATAAATGAAAGATCCCAGCAGCGCACGGAGATATTACATGAGCTACGATTCGCGCAAGATCCAATGATCATCAGTTTGCCTAGGCTCACTGACTCACACCCGCCAACCATACCGGCTATTTTGCATGCTTCTTTCGTTTCACCAACGATCTCGAATAATGTCCCATGACGCTTTATCATACACACAGAGCAGTGGCGCTACTGCATATGATTCGTTGATAGTTTTTTTTTTTTTATTCTGCATCTCCAAGCATTGCTCCTTATCACACGAAGATGCAAACTGAACGGCATTGCATCAGGGTCTGAGTTAGTGGCTCACTGTAAAGTGATGTGGGGGGTGGGGGGGGCTTCATTTGTTTATGGCACCTTTCGCTTTTTTGTGTCAGAGAGGCTTCATTTGTCTATGTCGCAGTATATTGCAAACACTCATTGCGATCATCTTCCAATTACGTCACATGGAAATGTTTGGTTTGAAGAGATCGCTATCAATGCAAAAACCTCCGTCTACAAACAGCTGACCATAAGGAACGGGCGGCTGTGTAAGGCAGCAACTTTTCAAGGGCCAAGATGCTCTCGATGCAGGAAAAGTGTAGACAAAACTGATAAGAACGCCCTCATATGCGAGTCGGTTCGAGAGCAATCAAGAGCTACACTTCCTGGGTTCGTCAGTTTGCTCCTAGCAGTGTGACACTAAACACTTCCCCGTGAGCGCTGTCCATAGTGTTTCCCTTGAAAA

5. **>AgSCRASP1**

CCAGATGCACGGGGATTAAACAATTCAAACAATAACCAAGACTCATAAAATCTTTATTTCCTATTCCTAATACTATAACGAGTCCGTTTATCCTAGCCTAGCTCTACCGCACGAATGTGTTTGCCCTCTCCCTTCGCTCCTCTGGTCAGCGACCCGGGAACTCACGCACATCCTTTCACGCACACATTATACGACAGCCGTCGACTGCTATCGGTGAGTGGTGCGCCCAGGTGGACAGAACTGCGTAGTCAGGTTGGACCACAATAGTATTCGTAATAGTGATGGGAAAAATGAAGATTTCGTCGGAATCGATTCCGGTTAGCTCCGAAGTTTTCTGGAATCGATTCCGGATAGTAGGTCCGGAATCAGTTTCCGGAATCGGCTCCGGAATCGGCTCCGGAATCGGAATCGTCTCCGGAATCGGAATCGGCTCCGGAATCGGAATCGGCTCTGGAATCGAATTTGGCTCCGGAATCAGATTCAACTCCGGAATCGGAATCGGGCTTCGGAATCAAAATCGGCTCCGACATCGGAATTGACTCCCAAATCAGAATTGGCTTCGAAACGGAGTCGGTTTCGGCATCTTCATAGGAATAGGCGTTTGGGATCAATGATGCTACTTGTTGATAGCGACAAAGAATCAAAATTTACTTGTATATGAATTCAAATGGAGATTGTTATTTCGCTTATTTATTATTTTCGCCCGATTGTTATTTCGCCCGGACTTATTTCGCTTCCCACACCTCAATTCAAAAACTAATTCACATTCCGGAGCTAACTCCATTTCTGGAGTCAAGCCCTACTGATTCCGTTACCGGCGCAAATTGCGTTTCCCAGGTCTCATCGCTACTTGCCTCCCATCTTAACGATCTTGCGCACCAACCAGTCAATTGAGCATTAATTGAACCCATTCTAGAAGCGTGCCCGACGGTGCTACCCCTTCTTCAATGAATCGGGGCATTTCACCGCATCATCGAATGTGTTTCACTGGAACCGCGTAGTGAAGTACCCTTCAGCGGGGAAAGTGCGACCGATTTGTAGCCATTTAATTGATTCATTAAATAGCGTCAATTTATTCGTCTACGCGGGCGCGCGCTCGCGCTATCATCTTGCGCCTCTTTCTATTCCGTACGCACGATGGCCATGATTCATCCCATTTTGTTCGGGTTTGTTTTGGGGGATTTACCACCGTGCGTGTGAGTGTGAAAGAAAAACATTGTCTCAACTGGGTAGTGAAGAGTTCTTAACGACGACACAGAGCCGTCGTAAGTAATCAACGCAAAGATGCTGATAAGAATGAAGAGAGAGAGCGAGAGACGCATCTAGCCGATGCGAGCGACGACGAACTGAGCTGGAAATTAAACACGAAGCGCGTCCGACTGACACTAAAACTGCCCGATATTCACTGGCTCTTTCATGCCGGTATCAACATATCGATAATAGGAGGGACGTGATTTTTTGCGAGAACGGTGGTGTAATTTCAAACACCTACCAAATGGTGTGTGAACCTTTATGGATCGTTATTCAAACCCGCGACACTTGGTGAAGGTGTTACGCCCGCTCGTGTTCTCGCTCAGGTCATTGAATAGCACCATGGTGTAATATTTAGAGCATCTTTGAGTAGTTGAGCATGAAATGAAAATGAAATAATGAATCTTAAAATGCATATGACATTGGTTATGGCTTTCCCGCACTAAATGACCACTTGGCACGATTAGTGTTGATGCGTCCATTCACTACACGAGTGACTCTTCATTCAAGCCGAACCAATTCCGTTGGTTCTTCGCGCCTTCTTCCAATTCCTTCGCGATCGTCGCTTCCTGTTGTTGTGAGCTCGCCCGAACCGAAGCGACTGACCTTTCGCTCTTCCCTTCCCGAGGGTGGAAGCATCTCTCCTAATCGGCACCAACTGATGATCGCTAGCCCAAGTCTGAAACGAGGGTCGCGGTCACACGACTGCTCATTGTACGCTCGTAACTGGACCAAGCAGGACGTCCAGTATCAGTGGCTCTTCGTTTAGTGATCTTGTTATTTTTTTTTTTTTGTGAGAGAATATTGTGCCATCACAGCCGCACCAGCTCGTGTCTGTGGTTTTGTGTATCTCCAGACAAAGATTAACAGTGACGGGATTAAAAGATAAGACACGAAGGAACCAGCTTCGAACGATCTTTAGTTGGGAGATTGGCGCAAGCGAAGATTTGCGCGAAAGCG
